# STIM1 and Endoplasmic Reticulum-Plasma Membrane Contact Sites Oscillate Independently of Calcium-Induced Calcium Release

**DOI:** 10.1101/2025.03.16.643575

**Authors:** Ding Xiong, Cheesan Tong, Yang Yang, Jeffery Yong, Min Wu

## Abstract

Calcium (Ca²⁺) release from intracellular stores, Ca²⁺ entry across the plasma membrane, and their coordination via store-operated Ca²⁺ entry (SOCE) are critical for receptor-activated Ca²⁺ oscillations. However, the precise mechanism of Ca²⁺ oscillations and whether their control loop resides at the plasma membrane or intracellularly remain unresolved. By examining the dynamics of stromal interaction molecule 1 (STIM1)—an endoplasmic reticulum (ER)-localized Ca²⁺ sensor that activates the Orai1 channel on the plasma membrane for SOCE—and in mast cells, we found that a significant proportion of cells exhibited STIM1 oscillations with the same periodicity as Ca²⁺ oscillations. These cortical oscillations, occurring in the cell’s cortical region and shared with ER-plasma membrane (ER-PM) contact sites proteins, were only detectable using total internal reflection fluorescence microscopy (TIRFM). Notably, STIM1 oscillations could occur independently of Ca²⁺ oscillations. Simultaneous imaging of cytoplasmic Ca²⁺ and ER Ca²⁺ with SEPIA-ER revealed that receptor activation does not deplete ER Ca²⁺, whereas receptor activation without extracellular Ca²⁺ influx induces cyclic ER Ca²⁺ depletion. However, under such nonphysiological conditions, cyclic ER Ca²⁺ oscillations lead to sustained STIM1 recruitment, indicating that oscillatory Ca²⁺ release is neither necessary nor sufficient for STIM1 oscillations. Using optogenetic tools to manipulate ER-PM contact site dynamics, we found that persistent ER-PM contact sites reduced the amplitude of Ca²⁺ oscillations without alteration of oscillation frequency. Together, these findings suggest an active cortical mechanism governs the rapid dissociation of ER-PM contact sites, thereby control amplitude of oscillatory Ca²⁺ dynamics during receptor-induced Ca²⁺ oscillations.

## Introduction

Ca²⁺ oscillations are a ubiquitous signaling mechanism observed across almost all cell types (Lewis and Cahalan, 1989; Millard et al., 1988; Rapp and Berridge, 1981; Woods et al., 1986), regulating processes such as actin dynamics, secretion (Tse et al., 1993; Wollman and Meyer, 2012), gene expression (Capite et al., 2009; Dolmetsch et al., 1998; Li et al., 1998), and arguably additional teleological functions (Tsien and Tsien, 1990). Since their discovery, two types of non-mutually exclusive models have emerged to explain their origin: one based on “cytoplasmic oscillator”, that focuses on oscillatory store release or Ca²⁺-induced Ca²⁺ release (CICR), and another involving “membrane oscillator”, that focuses on receptor-controlled periodic production of second messengers, such as inositol trisphosphate (IP₃), from the plasma membrane (Berridge et al., 1988; Berridge and Rapp, 1979). Despite extensive research, the mechanisms behind its oscillatory dynamics and the main site of control remain a matter of debate.

The CICR model proposes that Ca²⁺ oscillations result from periodic Ca²⁺ release from intracellular stores, primarily the ER, via an autocatalytic positive feedback loop (Goldbeter et al., 1990; Kuba and Takeshita, 1981). Experimental evidence for this model originated from excitable muscle cells (ENDO et al., 1970). Further support came from observations of Ca²⁺ oscillations persisting without extracellular Ca²⁺ entry (Wakui et al., 1990). Although such oscillations are typically short-lived in many systems, suggesting Ca²⁺ entry across the plasma membrane plays a significant role in either initiating or sustaining Ca²⁺ oscillations, Ca²⁺ oscillations in the absence of entry can be prolonged if Ca²⁺ extrusion is inhibited (Bird and Putney, 2005; Capite et al., 2009). This contributes to the prevailing view that Ca²⁺ release from the store (and the following refill) is the primary driver mechanism while Ca²⁺ entry primarily refills ER stores to sustain oscillatory release (Parekh, 2007).

An alternative model hypothesizes that Ca²⁺ oscillations could arise from membrane-dependent oscillators driving periodic second messenger production. IP₃, generated at the plasma membrane in response to receptor activation, binds to IP₃ receptor on the ER and regulates Ca²⁺ release from ER stores, has been proposed as such a messenger (Bezprozvanny et al., 1991; Ehrlich and Watras, 1988; Meyer et al., 1988; Meyer and Stryer, 1988). IP₃ oscillations have been observed in various systems (Harootunian et al., 1991; Hirose et al., 1999). However, while IP₃ can exhibit oscillatory dynamics, IP₃ oscillations typically occur alongside Ca²⁺ oscillations. Conversely, Ca²⁺ oscillations have been observed without IP₃ oscillations, such as when IP₃ levels are fixed (Wakui et al., 1989). Thus, it has been argued that IP₃ oscillations (or oscillations of related second messengers) may couple with Ca²⁺ oscillations but do not cause them. Despite extensive research into feedback between Ca²⁺ and IP₃, as well as observations of oscillations of other messengers like phosphatidylinositol 4,5-bisphosphate (PIP₂) (Wu et al., 2013; Xiong et al., 2016), and diacylglycerol (DAG)(Oancea and Meyer, 1998), no second messengers oscillations related to Ca²⁺ mobilization have been shown to occur independently of Ca²⁺ oscillations. This raises the question of whether an autonomous oscillatory process at the plasma membrane could control Ca²⁺ oscillations.

Because IP₃-Ca²⁺ oscillations are closely intertwined with Ca²⁺ release from ER stores, both CICR and IP₃-induced Ca²⁺ release models share the same prediction that the rate of ER store refilling determines the delay between successive Ca²⁺ transients and thus the oscillation period (Berridge, 2007, 1993; Goldbeter et al., 1990; Meyer and Stryer, 1988; Sneyd et al., 2004). However, this relationship has not been directly demonstrated in receptor-activated Ca²⁺ oscillations in non-excitable cells (Shuttleworth and Thompson, 1996). Studies on receptor-activated Ca²⁺ oscillations have not simultaneously monitored ER and cytoplasmic Ca²⁺ levels, leaving uncertainty about whether ER store refilling indeed acts as the rate-limiting step controlling oscillatory behavior.

In addition to IP₃ that originates from plasma membrane to regulate ER Ca²⁺ release, state of ER store release can be transduced back to Ca²⁺ entry through processes such as SOCE (Putney, 1986), a process first identified in mast cells and T cells (Hoth and Penner, 1992; Zweifach and Lewis, 1993). STIM1, a transmembrane protein residing in the ER, was identified as a key SOCE component that senses Ca²⁺ store depletion in the lumen of ER, translocates to ER-PM contact site, where it binds and activates Orai1, a plasma membrane calcium channel that mediates Ca²⁺ entry (Liou et al., 2005; Luik et al., 2008; Roos et al., 2005). In this study, we investigated the regulation of cortical STIM1 dynamics during receptor-mediated Ca²⁺ oscillations using antigen-stimulated RBL-2H3 mast cells as a model. Although SOCE is well-characterized, most studies of STIM1 recruitment have used nonphysiological stimuli such as SERCA ATPase inhibitor thapsigargin that completely deplete ER Ca²⁺. During receptor-activated Ca²⁺ oscillations, we observed cyclic STIM1 translocation, coupled with oscillations of other ER proteins at ER-PM contact sites. Co-imaging of STIM1 with cytoplasmic and ER-specific Ca²⁺ sensors revealed that STIM1 oscillations occur independently of Ca²⁺ oscillations or ER Ca²⁺ depletion. Furthermore, using optogenetic tools to stabilize ER-PM contact sites, we demonstrated that dynamic control of these sites modulates the amplitude of Ca²⁺ oscillations. These findings suggest an autonomous oscillatory process exist at the plasma membrane that is upstream of Ca²⁺ oscillations, and an underappreciated role of reversible ER-PM contact site interactions in controlling Ca²⁺ oscillations.

## Results

### STIM1 and Cortical ER-PM Contact Site Oscillations in Antigen-Stimulated Mast Cells

We studied the dynamics of STIM1 at the plasma membrane in mast cells (RBL-2H3) under antigen stimulation using total internal reflection fluorescence microscopy (TIRFM) (**Fig. 1a**). After sensitizing RBL cells with anti-DNP IgE and stimulating them with multivalent antigen DNP-BSA that crosslinks anti-DNP IgE, we observed an increase in STIM1 intensity at the ER-PM junction, followed by oscillatory behavior (**Fig. 1b, Supplemental Video 1**). These oscillations were characterized by the cyclic appearance of STIM1 puncta or fluctuations of these cortical signals near the plasma membrane (**Fig. 1c**), as no cyclic changes in fluorescence intensity were observed at higher z-planes using confocal microscopy (**Fig. 1—Supplemental Fig. 1**).

**Figure 1.**
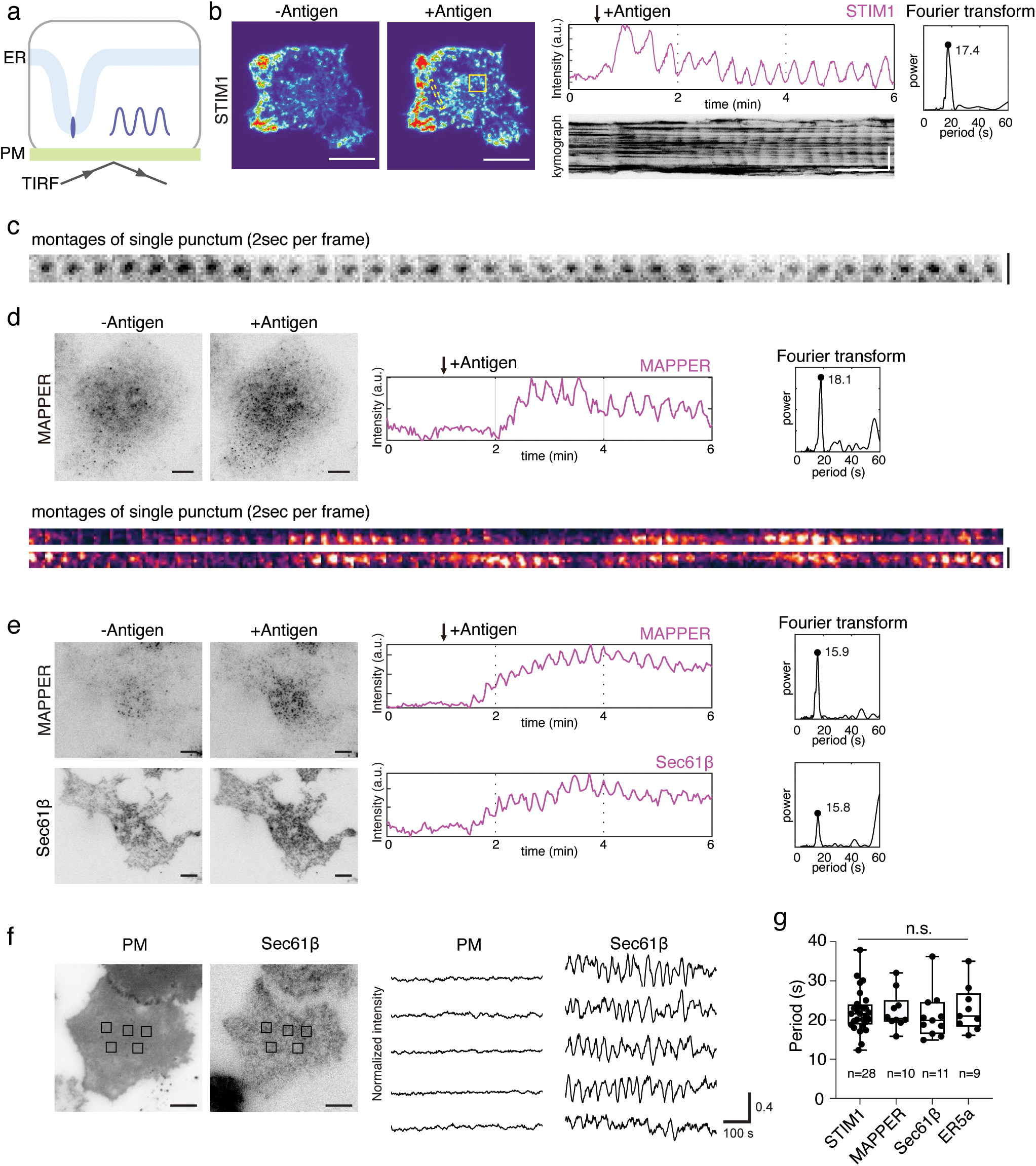
STIM1 and ER-PM contact site oscillations in antigen-stimulated RBL-2H3 mast cells. (**a**) Schematic of monitoring STIM1 and ER markers oscillations by TIRF microscopy. (**b**) Cyclic STIM1 oscillations upon antigen stimulation. *Left*: Representative TIRF images of cell expressing STIM1-RFP before (-Antigen) and after (+Antigen) DNP-BSA stimulation (n= 36 cells from 12 experiments). Images are shown with “thermal” lookup table. *Middle*: Representative STIM1 intensity profile and kymograph for STIM1 oscillations. *Right*: Period of STIM1 oscillations as analyzed by Fourier transform, with the period of major peak labelled with black dot. (**c**) Montage of single punctum of STIM1-RFP in 35 consecutive frames (2-s interval). (**d**) Representative TIRF images of GFP-MAPPER before and after DNP-BSA stimulation, with intensity profile, Fourier transform of intensity profile, and sequential montage of single punctum of 60 frames (2-s interval). (**e**) Representative TIRF images, intensity profile and Fourier transform of GFP-MAPPER and mCherry-Sec61β before and after DNP-BSA stimulation (n=10 cells from 3 experiments). (**f**) Representative TIRF images and intensity profiles of cell expressing PM-GFP and mCherry-Sec61β quantified in multiple regions of interests (n=12 cells from 3 experiments). (**g**) Box and whisker plots showing the periodicities of STIM1 and ER markers oscillations. Statistics analyzed by Brown-Forsythe and Welch ANOVA test, n.s., not significant. For TIRF images, scale bar=10 µm; for kymographs, scale bar=1 min (horizontal bar), 10 µm (vertical bar); for montage, scale bar=2 µm.

To determine whether STIM1 oscillations indicate increased ER-PM contact site formation, we examined the dynamics of ER-PM contact site marker MAPPER (Chang et al., 2013). We could observe oscillations of MAPPER which was contributed by increased MAPPER puncta formation as well as oscillations of individual puncta (**Fig. 1d, Supplemental Video 2**). Observation of these MAPPER oscillations require minimal expression level of the MAPPER sensor, as cells with high expression level did not show oscillations (**Fig. 1—Supplemental Fig. 2**). Interestingly, additional ER markers, such as Sec61β and ER5a-GFP could also oscillate even though they did not show clear puncta at membrane contact, suggesting increased signals of cortical ER in the TIRF field could contribute to oscillations (**Fig. 1e, Supplemental Video 3**). We also ruled out the possibility that these oscillations were caused by plasma membrane fluctuations, as plasma membrane markers like PM-GFP did not oscillate in the same cell when cortical ER oscillations occurred (**Fig. 1f**). All ER markers show similar periods in their oscillations in the range of 22±5 sec (**Fig. 1g**). When comparing the amplitude of oscillations for STIM1 and the ER marker ER5a-GFP, we found that ER5a exhibited oscillations with amplitudes comparable to those of STIM1 (**Fig. 1—Supplemental Fig. 3a-b**). However, under non-physiological conditions, such as ER calcium store depletion induced by the thapsigargin, STIM1 recruitment to the ER-PM junction was elevated compared to the ER5a signal (**Fig. 1—Supplemental Fig. 3c-d**). These results suggest that ER-PM junctions can oscillate under physiological stimulation and that these oscillations are mainly contributed by dynamic translocations of cortical ER and oscillatory assembly of ER-PM contact sites.

### STIM1 Oscillations Can Occur Independently of Cytoplasmic Ca²⁺ Oscillations

We next examined the correlation between STIM1 oscillations and calcium oscillations. To monitor cytoplasmic calcium levels, we used the genetically encoded calcium indicator GCaMP3. Surprisingly, STIM1 oscillations were not always coupled with calcium oscillations but instead exhibited a more complex relationship. To quantify these heterogeneous responses, we categorized our data into three groups based on patterns of calcium responses. While there is a general trend of stronger calcium oscillations immediately after stimulation, followed by a gradual decline over time, there are significant variabilities within a cell population, both within and across experiments. Some cells exhibit short-lived calcium oscillations, whereas others in the same sample display persistent oscillatory activity. Therefore, classifying single-cell data solely based on time after stimulation is unreliable (**Fig. 2a**). Instead, we analyzed calcium traces with a ten-minute sampling window by applying Fourier transform analysis to converting intensity profiles from the time domain to the frequency domain. The first group consisted of traces with persistent calcium oscillations. When converted in the Fourier space, they displayed a single prominent peak, confirming their periodic nature (**Fig. 2b**). The second group contains traces with intermittent calcium spikes but no distinct peaks in the Fourier space (**Fig. 2c**). The third group showed no calcium spikes within the ten-minute window (**Fig. 2d**).

**Figure 2.**
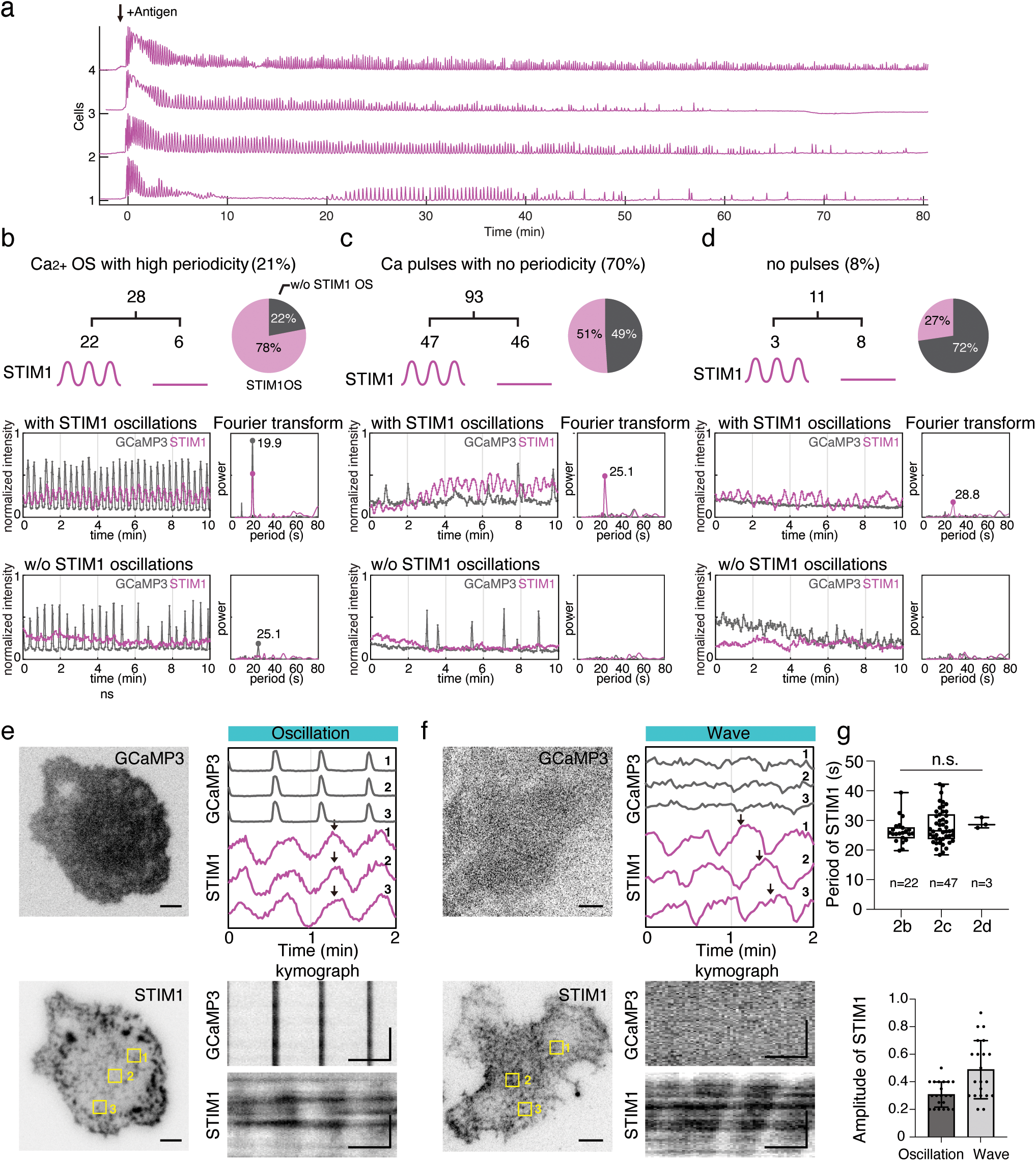
STIM1 oscillations can occur independently of cytoplasmic Ca^2+^ oscillations. (**a)** Representative profiles reveal variations in Ca^2+^ signal duration upon antigen stimulation in RBL-2H3 cells expressing GCaMP3. **(b-d**) Profiles of Ca^2+^ (GCaMP3) and STIM1-RFP classified using a sampling window of 10 min acquired at 2s per frame (n=132 cells from 4 experiments). *Upper*: Summary of the percentage of the cells in each category. *Bottom*: Intensity profiles and Fourier analysis of STIM1-RFP and GCaMP3 fluorescence in each category. Data are 10-40min after antigen stimulation. (**e-f**) Representative images, intensity profiles and kymographs of cells with STIM1 oscillations coupled with Ca^2+^ oscillations (e) or uncoupled with Ca^2+^ oscillations (f). Three regions of interest were used for every cell. (**g**) Box and whisker plots showing the period of STIM1 oscillations in each category. Statistics analyzed by Brown-Forsythe and Welch ANOVA test, n.s., not significant. (**h**) Statistics for the amplitude of STIM1 oscillation and wave (n=21 and 20 cells, respectively). For TIRF images and montage, scale bar=5 µm; for kymographs, scale bar= 30s (horizontal bar), 5 µm (vertical bar).

Among the cells with persistent calcium oscillations (21%; 28 out of 132 cells), most (89%) displayed coupled STIM1 oscillations (**Fig. 2b, Supplemental Video 4**), indicating a strong prevalence of STIM1 oscillations during calcium oscillations. In these coupled events, STIM1 and calcium oscillations showed matching frequencies (**Fig. 2b**). Conversely, in cells exhibiting calcium oscillations without coupled STIM1 oscillations, the Fourier transform revealed a major peak for calcium but not for STIM1 (**Fig. 2b**). In the second group, comprising cells with irregular calcium spikes (70%; 93 out of 132 cells), 62% displayed STIM1 oscillations (**Fig. 2c, Supplemental Video 4**). However, the cyclic patterns of STIM1 oscillations in these cells were not coupled with calcium spikes (Fig. 2c). In the third group, consisting of cells with no calcium spikes (8%; 11 out of 132 cells) (**Fig. 2d, Supplemental Video 4**), 36% exhibited STIM1 oscillations (**Fig. 2d**).

Upon closer examination of the cyclic STIM1 translocation patterns, we observed distinct spatial patterns between conditions with and without calcium oscillations. When calcium oscillations were present, STIM1 translocation occurred synchronously across different regions of the same cell (**Fig. 2e, Supplemental Video 4**). In contrast, in the absence of calcium oscillations, STIM1 puncta translocation was asynchronous across different regions of the cell, manifesting as traveling waves (**Fig. 2f, Supplemental Video 4**). The STIM1 oscillations coupled with calcium oscillations have shorter period and lower amplitude than that of uncoupled with calcium oscillations (**Fig. 2g**).

The presence of STIM1 oscillations in cells lacking cytoplasmic calcium oscillations was unexpected. We observed interconversion between calcium oscillation-coupled and uncoupled STIM1 oscillations. For instance, STIM1 oscillations could persist after calcium oscillations stopped (**Fig. 2—Supplemental Fig. 1a**). Additionally, cells transitioning from sustained to intermittent calcium oscillations maintained cyclic STIM1 oscillations (**Fig. 2— Supplemental Fig. 1b**). These findings suggest that STIM1 oscillations coupled and uncoupled with calcium oscillations represent transient dynamical states, rather than being due to intrinsic cell-to-cell variability.

### STIM1 Oscillations Are Not Driven by Oscillatory ER Ca²⁺ Depletion

The observation of STIM1 oscillations in the absence of cytoplasmic Ca²⁺ oscillations suggests that STIM1 oscillations may not simply happen as the result of cytoplasmic Ca²⁺ oscillations. Given that STIM1 serves as a sensor for ER Ca²⁺, we next examined whether ER Ca²⁺ depletion occur in a cyclic fashion and whether STIM1 oscillations could be driven by oscillatory ER Ca²⁺ depletion.

To investigate whether ER Ca²⁺ depletion coincided with cytoplasmic Ca²⁺ oscillations, we simultaneously monitored ER Ca²⁺ levels using biosensor CEPIA-ER (Suzuki et al., 2014) and cytosolic Ca²⁺ levels using GCaMP3 in the same cell via TIRFM (**Fig. 3a**). To assess how CEPIA-ER respond to a broad dynamic range of ER Ca²⁺ levels, we first stimulated RBL cells with DNP-BSA antigen in a Ca²⁺-free buffer to deplete ER stores, then introduced a Ca²⁺-containing buffer to restore ER Ca²⁺ levels, and finally re-depleted ER Ca²⁺ again using thapsigargin.

**Figure 3.**
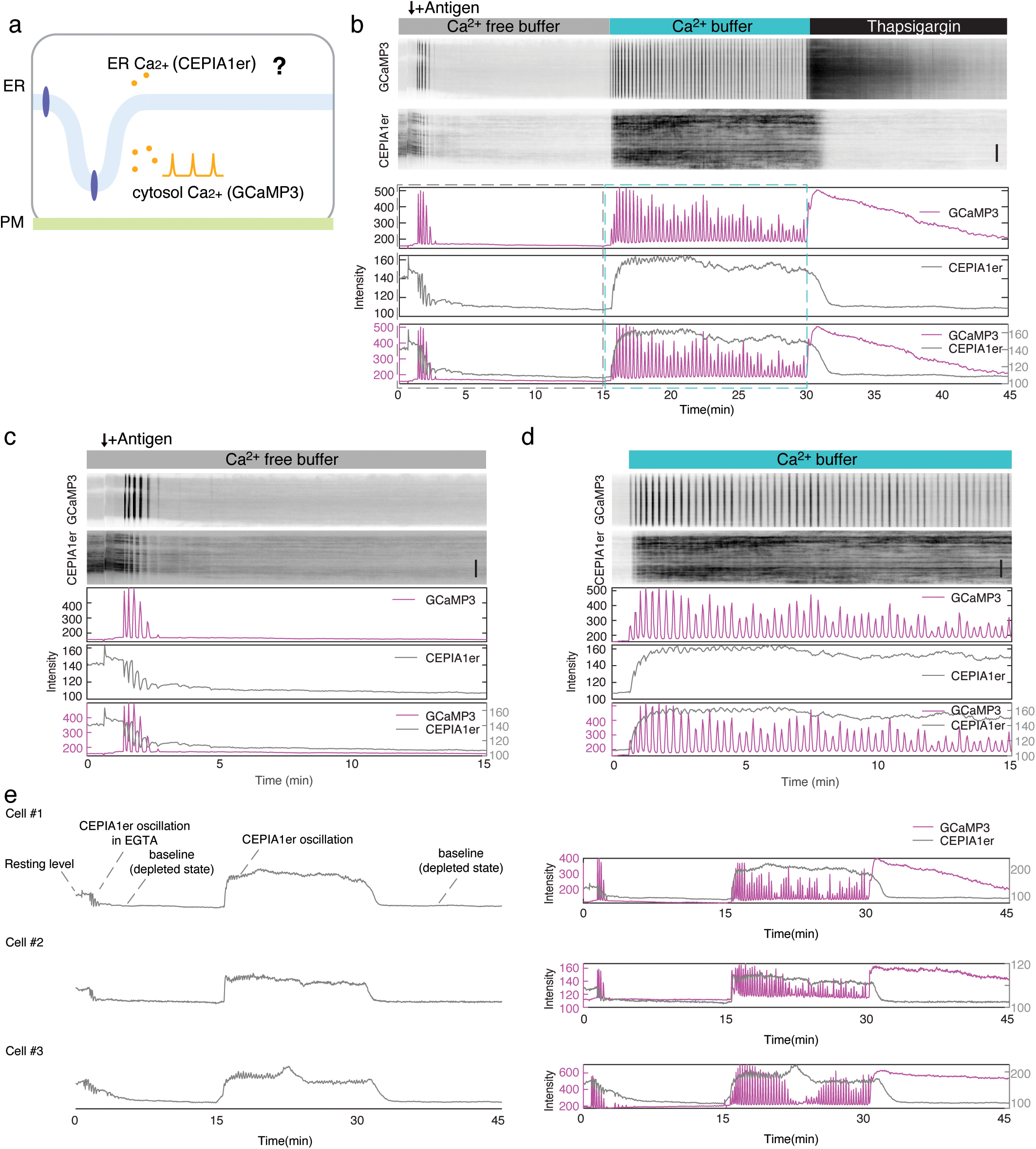
Simultaneous visualization of ER and cytoplasmic Ca^2+^ oscillations showing ER Ca^2+^ depletion only occurs in non-physiological conditions. **(a)** Schematic of cytoplasmic and ER Ca^2+^ sensor used. **(b)** Representative kymograph and intensity profiles of GCaMP3 and CEPIA1er after DNP-BSA stimulation in Ca^2+^ free buffer, followed by replacement with Ca^2+^ buffer and the final addition of thapsigargin. Dashed line indicate time period shown in (c-d) **(c-d)** Zoomed-in of (b) in the Ca^2+^ free and Ca^2+^ phases. **(e)** Intensity profiles of GCaMP3 and CEPIA1er from additional examples, as stimulated in (b). Scale Bar=10 µm.

In the first phase, stimulation of cells with DNP-BSA antigen in the Ca²⁺-free buffer (with EGTA) triggered Ca²⁺ release solely from the internal ER Ca²⁺ pool, without Ca²⁺ influx from the extracellular space. This elicited transient Ca²⁺ oscillations, lasting 2.9 ± 1.2 cycles (148 cells across six experiments) (**Fig. 3—Supplemental Fig. 1, Supplemental Video 5**). Under these conditions, ER Ca²⁺ levels exhibited anti-coupling with cytosolic Ca²⁺ oscillations (**Fig. 3b**). Each Ca²⁺ spike correlated with pulses of CERPIA-ER depletion, with decreasing baseline, ultimately reaching a flat baseline when ER Ca²⁺ level insufficient to sustain further cytosolic Ca²⁺ oscillations (**Fig. 3c**). In the second phase, switching to a Ca²⁺-containing buffer restored robust cytoplasmic Ca²⁺ oscillations (**Fig. 3b**). The intensity of CEPIA-ER signals increased and displayed low level oscillations that were coupled with some, but not all, cytoplasmic Ca²⁺ oscillations (**Fig. 3d**). In the third phase, the addition of the SERCA ATPase inhibitor thapsigargin caused an acute drop in CEPIA-ER signals to the same baseline as in the EGTA phase, while cytoplasmic Ca²⁺ levels increased, confirming Ca²⁺ mobilization from the ER to the cytosol (**Fig. 3b**). Collectively, the overall trend in the CEPIA-ER dynamics —decrease with EGTA, recovery to an elevated amplitude in physiological buffer, and depletion with thapsigargin—aligns with the expected changes in ER Ca²⁺, indicating that CEPIA-ER can sense ER Ca²⁺ changes.

Notably, during the second phase of cytoplasmic Ca²⁺ oscillations, the CEPIA-ER signal increased beyond the resting level prior to stimulation observed in the first phase (**Fig. 3e**). This suggests that, rather than being depleted, ER Ca²⁺ stores were replenished to levels above resting level following receptor activation. Low-amplitude oscillations in CEPIA-ER signals were observed; however, the lowest points of these oscillations remained higher than the initial resting level before stimulation and significantly above the baseline corresponding to ER Ca²⁺ store depletion. The lack of ER Ca²⁺ depletion is also unlikely to be due to the slow dissociation rate of the Ca²⁺ sensor, as ER Ca²⁺ depletion was clearly detectable by CEPIA-ER in the first phase when extracellular Ca²⁺ was absent. In addition, it is unlikely that the baseline in the second phase was elevated due to changes in cell shape because thapsigargin treatment returned CEPIA-ER signals to the same baseline as in the EGTA phase. Therefore, we conclude that ER Ca²⁺ levels were minimally depleted during receptor-activated Ca²⁺ oscillations.

Importantly, while CEPIA-ER oscillations were clearly observed in both the EGTA phase and some of the persistent Ca²⁺ oscillation phase, their dynamics did not follow the predictions of CICR models, in which the rate of ER Ca²⁺ store recovery determines oscillation periodicity. When zooming in on individual spikes, we found that ER Ca²⁺ depletion was replenished by the time each cytosolic Ca²⁺ spike ended and CEPIA-ER signals exhibited a plateau phase during the interspike period of cytoplasmic Ca²⁺ oscillations (**Fig. 4a**, **Fig. 4—Supplemental Fig. 1**), indicating that store refilling was not rate-limiting for the subsequent Ca²⁺ spike.

**Figure 4.**
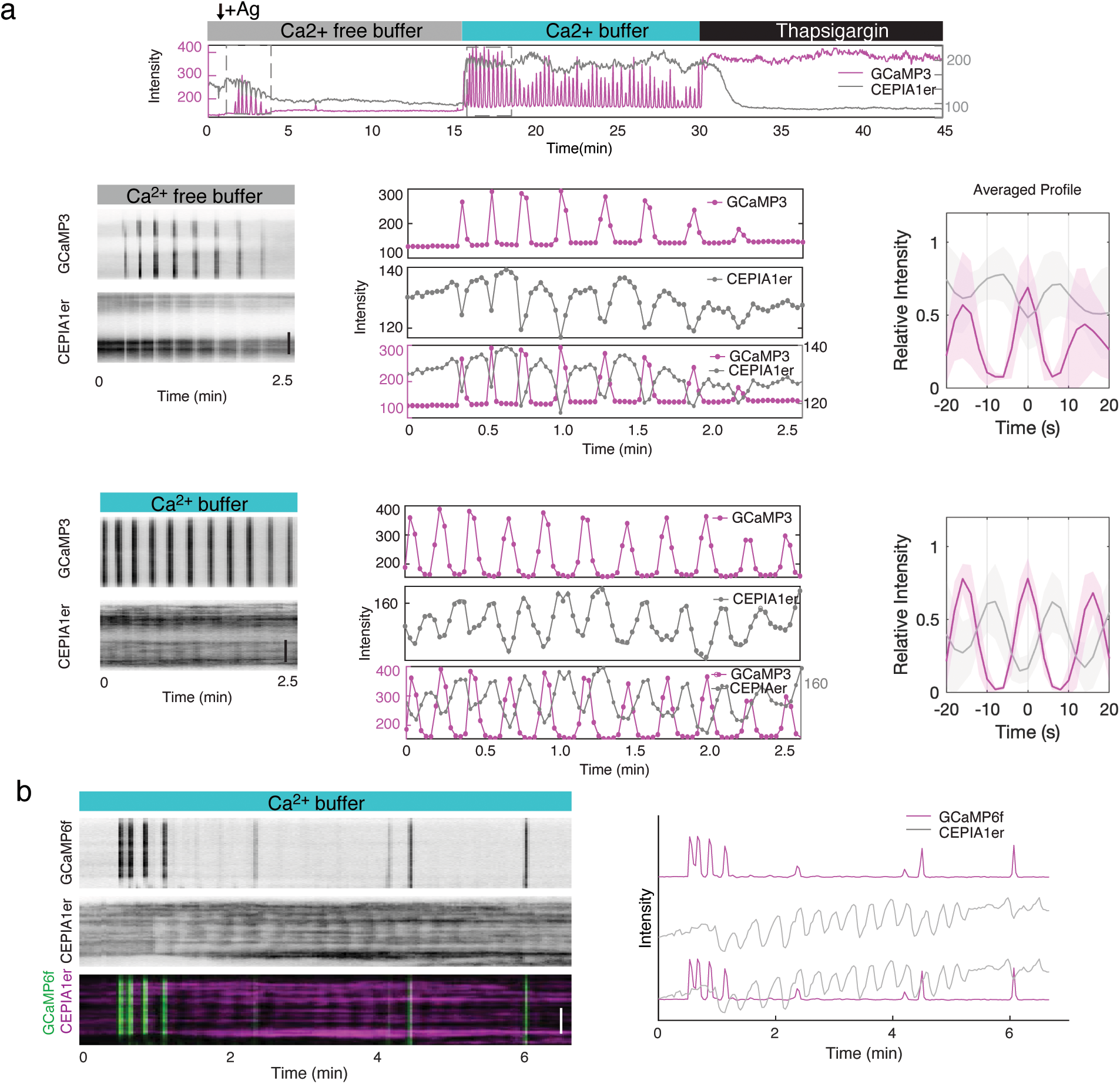
Ca^2+^ store refilling was not rate-limiting for the subsequent cytosplasmic Ca²⁺ spike. **(a)** Intensity profile, zoomed-in kymograph and high magnification view of intensity profiles of GCaMP3 and CEPIA1er after DNP-BSA stimulation in Ca^2+^ free buffer, followed by replacement with Ca^2+^ buffer and the final addition of thapsigargin. **(b)** Examples of CEPIA1er oscillations that are uncoupled with GCaMP3 oscillations and the corresponding kymograph showing CEPIA1er oscillations in this context were not synchronized across the cell. Scale Bar=10 µm.

One potential caveat of the CEPIA-ER probe is that, since it is localized on the ER, its fluctuating signal may partially reflect ER-PM dynamics. Indeed, we observed CEPIA-ER oscillations that were not coupled with cytoplasmic Ca²⁺ oscillations (**Fig. 4b**). These CEPIA-ER oscillations that were not coupled with cytoplasmic Ca²⁺ oscillations could be distinguished from those in the EGTA phase based on their spatial patterns in kymographs. CEPIA-ER oscillations in the EGTA phase were anti-correlated with cytoplasmic Ca²⁺, synchronized across the cell and appeared as vertical lines in the kymographs (compare **Fig. 4b** with **Fig. 4a**, EGTA phase). Additionally, when imaging was performed under non-TIRF (HiLo) conditions that could not detect cortical ER oscillations, synchronized CEPIA-ER oscillations persisted (**Fig. 4—Supplemental Fig. 2**). This suggests that the synchronized depletion in CEPIA-ER signals reports ER Ca²⁺ dynamics. We therefore conclude that CEPIA-ER oscillations in the EGTA phase reflect oscillatory release of ER Ca²⁺, whereas CEPIA-ER oscillations in other contexts are subjective to the compounding effect of ER movement and may or may not indicate ER Ca²⁺ depletion.

### Nonphysiological Oscillatory ER Ca²⁺ Release Trigger STIM1 Accumulation but Not Oscillations

The simultaneous imaging of ER Ca²⁺ with CEPIA-ER likely report an upper limit of the extent of ER depletion, which was significant under nonphysiological conditions (e.g., EGTA treatment). Under physiologically induced Ca²⁺ oscillations, ER Ca²⁺ levels were far from depleted, instead there is an overflow Ca²⁺ compared to the resting level. However, the possibility of transient ER Ca²⁺ depletion with the global ER Ca²⁺ overflow cannot be entirely ruled out. To test the impact of such mechanism on STIM1 dynamics, we examined whether ER Ca²⁺ release, when occurring out of phase with cytoplasmic Ca²⁺, could be sufficient to induce STIM1 oscillations. We performed three-color imaging to simultaneously monitor STIM1 dynamics, ER Ca²⁺ (via CEPIA-ER), and cytosolic Ca²⁺ (via GCaMP3) in cells stimulated in a Ca²⁺-free buffer (**Fig. 5a**). Under these conditions, each cytoplasmic Ca²⁺ spike was coupled with an acute decrease in ER Ca²⁺, as evidenced by the clear out-of-phase vertical stripes in the kymographs (**Fig. 5a**). Consistent with STIM1 sensing ER Ca²⁺ depletion, STIM1 intensity increased significantly. However, this increase did not result in an oscillation; instead, it exhibited a continuous elevation over time (**Fig. 5a-b**). This suggests that when STIM1 responds to oscillatory ER store depletion, its accumulation is not rapidly reversible on the timescale of these Ca²⁺ oscillations. Together, these results indicate that not only are ER Ca²⁺ oscillations unlikely to occur under conditions of receptor-induced Ca²⁺ oscillations, but even if they do, ER Ca²⁺ oscillations alone are not sufficient to drive STIM1 oscillations within the observed timescale of 15–30 seconds.

**Figure 5.**
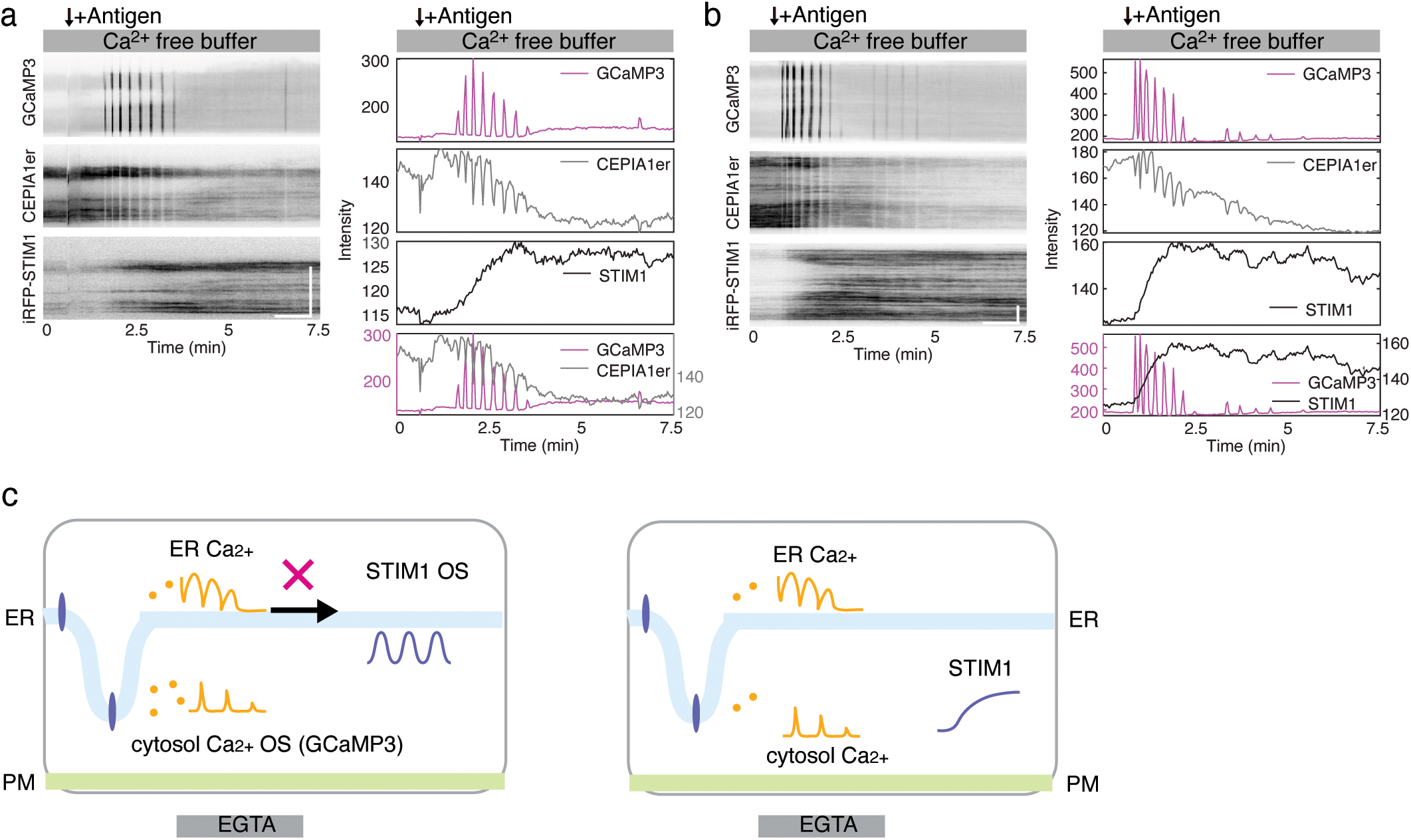
Non-physiological oscillatory ER Ca²⁺ release trigger STIM1 accumulation but not oscillations. (**a-b**) Two representative kymograph and intensity profiles of GCaMP3 and CEPIA1er, stimulated with DNP-BSA in Ca^2+^ free buffer. (**c**) Schematic illustration depicting ER Ca²⁺ release trigger STIM1 accumulation but not oscillations. Scale Bar=10 µm.

### Persistent ER-PM Contact Sites Inhibit Ca²⁺ Oscillations and Modulate Their Amplitude

We have shown that STIM1 and the cortical ER oscillate on a timescale that matches the frequency of receptor-induced Ca²⁺ oscillations. Importantly, cortical ER reversibly associates with the plasma membrane, and its dissociation does not occur as a result of cytoplasmic or ER Ca²⁺ oscillations. To investigate the significance of ER-PM contact site dissociation for Ca²⁺ oscillations, we stabilized ER-PM contact using a series of optogenetic tools based on light-induced dimerization between Cry2 and CIBN (Xiong et al., 2016) and test their impact on cytoplasmic Ca²⁺ oscillations.

In the first method, we fused STIM1 to Cry2 (**Fig. 6a**). A 7-amino-acid linker (GILQSTM) and an unstructured 243-amino-acid stretch after the SOAR domain of STIM1 ensured flexibility for membrane binding upon light-induced translocation. Upon blue light activation, STIM1-Cry2-miRFP670 was rapidly recruited to the TIRF field as distinct puncta in cells transfected with the plasma membrane-localized CIBN-GFP-CAAX (**Fig. 6b**). We further validated STIM1-Cry2-miRFP670 functionality by examining its translocation upon thapsigargin treatment. Without light induction, STIM1-Cry2-miRFP670 formed distinct puncta, showing a significant intensity increase (115 ± 95%) upon thapsigargin treatment (**Fig. 6c**, **Fig. 6—Supplemental Video 6**). This confirmed that Cry2 did not disrupt STIM1’s function as an ER calcium sensor.

**Figure 6.**
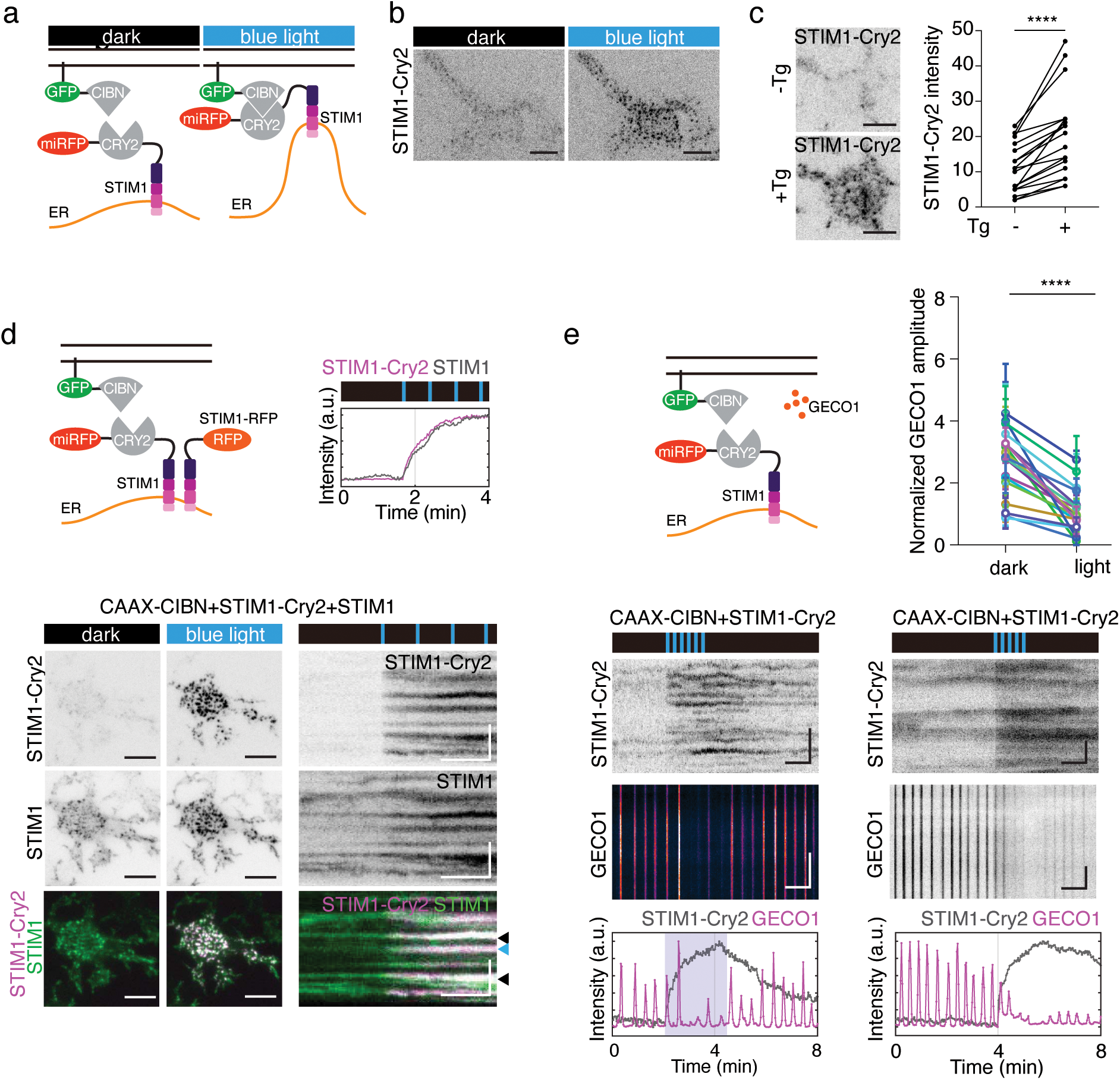
Optogenetic recruitment and stabilization of STIM1 to plasma membrane inhibits Ca²⁺ oscillations. (**a**) Schematic of light-induced plasma membrane translocation of STIM1-Cry2-miRFP670 by CIBN-GFP-CAAX. (**b**) Representative TIRF images of a cell before and after blue light exposure without antigen stimulation (n=24 cells from 9 experiments). (**c**) TIRFM images and quantification of STIM1-Cry2-miRFP670 intensity upon 10μM thapsigargin (Tg) treatment, without antigen stimulation (n=19 cells from 3 experiments), **** *P*<0.0001. (**d**) Schematic of experiment and intensity profile (*upper*), and TIRF images with kymographs (*bottom*) showing coupled translocation of STIM1-Cry2-miRFP670 and STIM1-RFP upon blue light exposure, without antigen stimulation (n=6 cells). Black and blue arrows show pre-existing or newly formed STIM1 puncta, respectively. (**e**) Schematic, kymographs, intensity profiles and quantification showing the effect of STIM1-Cry2-miRFP670 translocation on Ca^2+^ (GECO1) oscillations. Movies were acquired 10min after antigen stimulation. Cell showing decreased (*bottom left*) or almost abolished (*bottom right*) Ca^2+^ oscillations when STIM1-Cry2-miRFP670 is recruited upon blue light pulses. Pair-wise quantification for intensity of average spike amplitude in Ca^2+^oscillations before and after light pulses in different cells (n= 175, 167 peaks, respectively, 18 cells from 9 experiments), **** *P*<0.0001. Error bar, s.d. Figure 6c is quantified by paired two-tailed *t*-test and figure 6e is quantified by Welch’s *t*-test. For TIRF images, scale bar=10 µm; for kymographs, scale bar=1min (horizontal bar), 5 µm (vertical bar).

To test whether anchoring STIM1-Cry2-miRFP670 to the membrane interferes with the recruitment of other proteins, we assessed STIM1-RFP translocation. Following blue light activation, STIM1-RFP colocalized with STIM1-Cry2-miRFP670 puncta, indicating clustering of STIM1-Cry2-miRFP670 with STIM1-RFP. STIM1-RFP intensity increased significantly (40 ± 20%) alongside STIM1-Cry2-miRFP670, confirming that STIM1-Cry2-miRFP670 can recruit STIM1 to ER-PM contact sites (**Fig. 6d**, **Supplemental Video 7**). Most STIM1 puncta were preexisting before light induction and increased in intensity after STIM1-Cry2-miRFP670 translocation, while a few new puncta formed (**Fig. 6d**).

Next, we evaluated the effect of stabilized STIM1 translocation on Ca²⁺ oscillations. Sustained Ca²⁺ oscillations were decreased or inhibited in 69% of cells (18 of 26 cells, 9 experiments) (**Fig. 6e**). Among these, 15 cells showed a decrease in maximal GECO1 intensity, which recovered upon removal of blue light pulses. In 3 cells, Ca²⁺ oscillations were nearly completely inhibited. On average, Ca²⁺ oscillation amplitude was reduced by approximately 57 ± 19%.

In a second approach, we confirmed that the inhibitory effect was not dependent on the type of lipid anchor or the specific lipid domain targeted. To test this, we replaced the plasma membrane anchor for CIBN, switching from CAAX—a prenylation motif that targets proteins to the inner leaflet of the plasma membrane, predominantly in non-raft regions—to Lyn11, a saturated lipid modification from Lyn kinase that directs proteins specifically to lipid rafts, the more ordered, cholesterol- and sphingolipid-rich microdomains of the membrane. Previous work suggested that STIM1–Orai1 association could be sensitive to ordered or disordered lipid domains (Calloway et al., 2011; Maléth et al., 2014). We repeated the experiment using Lyn11-CIBN-GFP as the plasma membrane anchor to recruit STIM1-Cry2-miRFP670 and observed similar inhibitory effects (**Fig. 6—Supplemental Fig. 1a**). Likewise, in cells where STIM1-Cry2-miRFP670 was targeted to Lyn11-CIBN-GFP, Ca²⁺ oscillations were either abolished or persisted with reduced amplitudes (**Fig. 6—Supplemental Fig. 1b**).

Finally, in a third approach, we addressed concerns that overexpression of STIM1-Cry2-miRFP670 might disrupt the endogenous STIM-Orai complex stoichiometry or that the Cry2 tag on STIM1 might exert dominant-negative effects inhibiting Ca²⁺ oscillations. To bypass these potential issues, we fused the ER transmembrane domain of Sac1 (Sac1TM) to Cry2, allowing artificial ER-PM contact formation by recruiting Cry2-Sac1TM to the plasma membrane (**Fig. 7a**). To ensure flexibility for membrane binding upon light activation, we incorporated the same linker (GILQSTM) and an unstructured 243-amino-acid stretch following the Sac1 transmembrane domain. Upon blue light activation, Sac1TM-Cry2-miRFP670 rapidly formed distinct puncta within the TIRF field (**Fig. 7b-c**).

**Fig 7.**
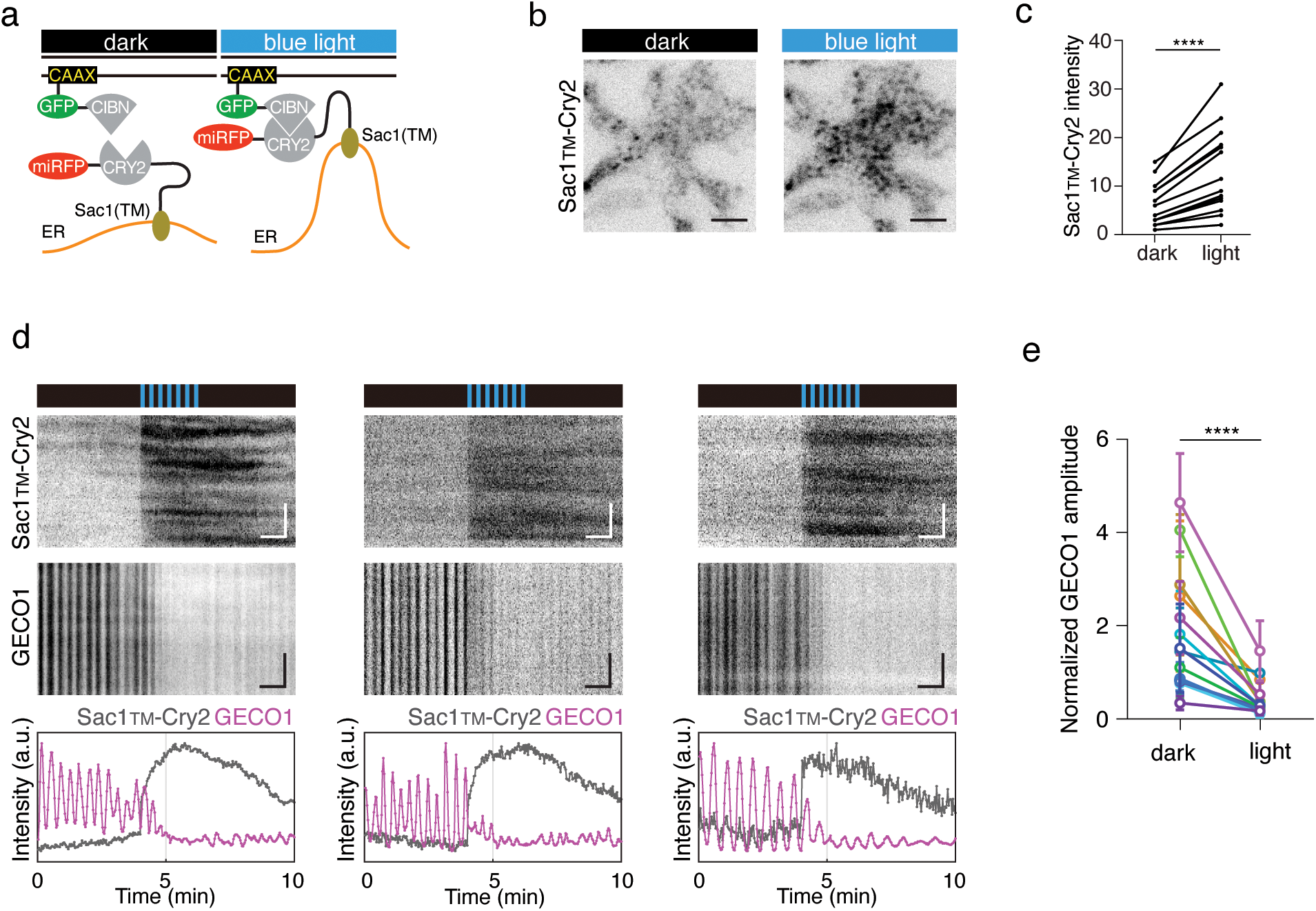
Optogenetic recruitment and stabilization of the transmembrane domain of Sac1 (Sac1_TM_) to plasma membrane inhibits Ca²⁺ oscillations. (**a**) Schematic of light-induced plasma membrane translocation of Sac1_TM_-Cry2-miRFP670 by CIBN-GFP-CAAX. (**b-c**) Representative TIRF images and quantification of Sac1_TM_-Cry2-miRFP670 intensity before and after blue light exposure without antigen stimulation (n=14 cells), **** *P*<0.0001. (**d**) Kymographs and intensity profiles of Sac1_TM_-Cry2-miRFP670 and GECO1 during exposure to blue light pulses. Movies were acquired 10 min after antigen stimulation. Cell showing decreased or almost abolished Ca^2+^ oscillations when Sac1_TM_-Cry2-miRFP670 is recruited. (**e**) Pair-wise quantification for average amplitude of spikes in Ca^2+^ oscillations before and after light pulses (n= 128, 160 peaks, respectively, 14 cells). Error bar, s.d. **** *P*<0.0001. Figure. 7c quantified by paired two-tailed *t*-test, Figure. 7e quantified by Welch’s *t*-test. For TIRF images, scale bar=10 µm; for kymographs, scale bar=1min (horizontal bar), 5 µm (vertical bar).

Stabilizing ER-PM contact with Sac1TM-Cry2 significantly impacted Ca²⁺ oscillations. In 93% of cells exhibiting sustained Ca²⁺ oscillations, blue light stimulation reduced oscillation amplitude or completely inhibited oscillations, indicating that persistent ER-PM contact suppressed Ca²⁺ dynamics (**Fig. 7d**). Among these, 57% of cells exhibited a decrease in maximal GECO1 intensity, while 36% showed near-complete inhibition of Ca²⁺ oscillations. On average, Ca²⁺ oscillation amplitude was reduced by approximately 75 ± 14% (**Fig. 7e**).

Remarkably, when Ca²⁺ oscillation amplitudes were reduced upon optogenetic recruitment of membrane contact sites—regardless of the dimerization strategy—their frequencies remained unchanged (**Fig. 8a**). We also observed cases where Ca²⁺ oscillations persisted after the initial optogenetic perturbation, allowing an additional round of STIM1-Cry2 recruitment, which further reduced oscillation amplitude while maintaining the same frequency (**Fig. 8b**). These results indicate that light-induced persistent ER-PM contact inhibits oscillatory Ca²⁺ responses and modulates their amplitudes, suggesting that the reversibility of ER translocation plays a role in tuning antigen-induced Ca²⁺ oscillations. While Ca²⁺ oscillations are well recognized as frequency-encoded signals, we are unaware of any previous experimental perturbation that selectively modulates oscillation amplitude without altering frequency.

**Fig 8.**
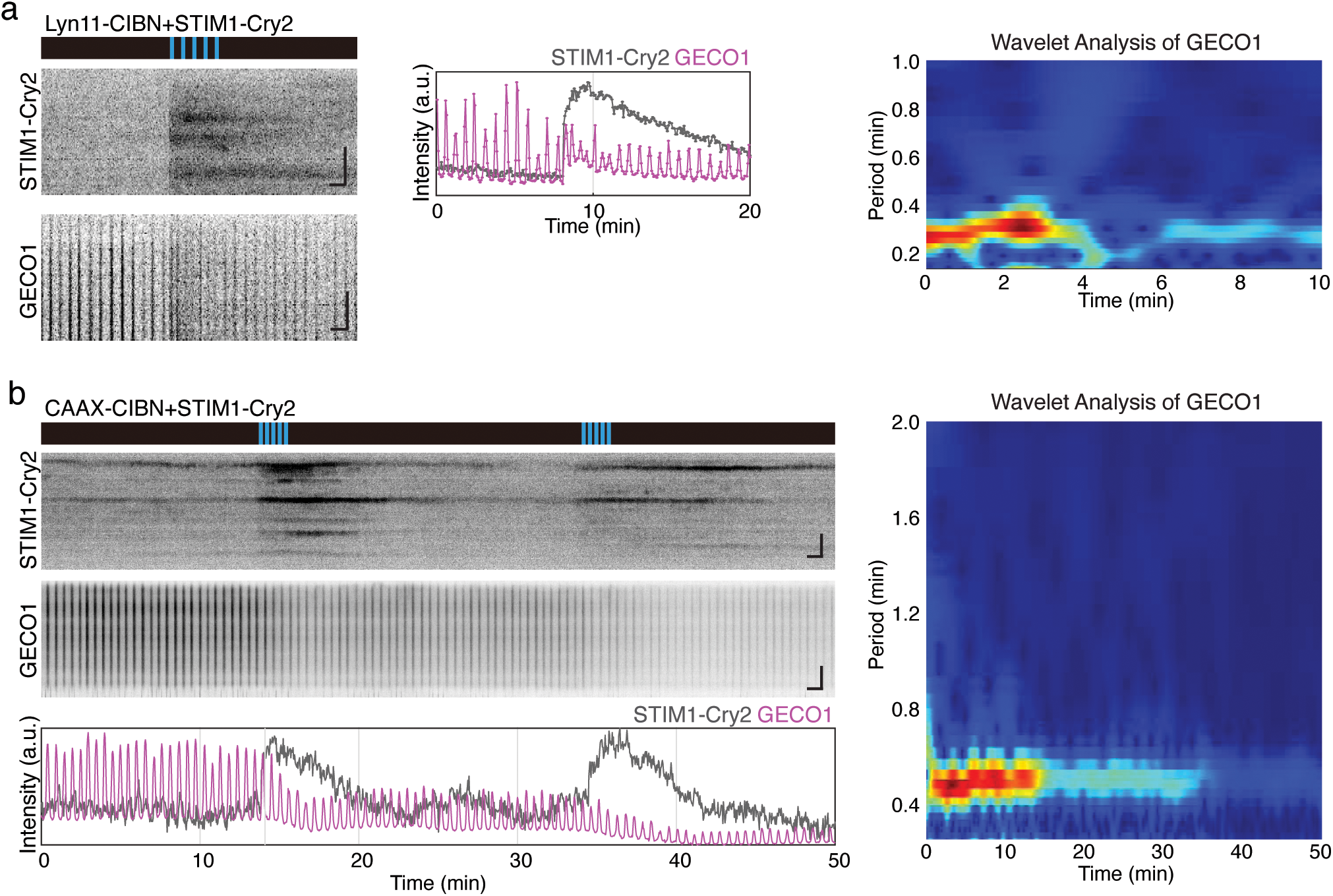
Uncoupling of Ca²⁺ oscillations amplitude and frequency by optogenetic perturbation of ER-PM contact. (**a**) Kymographs, intensity profile and wavelet analysis of a cell expressing STIM1-Cry2-miRFP670 and GECO1 during exposure to blue light pulses. (**b**) Kymographs, intensity profile and wavelet analysis of a cell expressing STIM1-Cry2-miRFP670 and GECO1 during two rounds of exposure to blue light pulses. Scale bar=1min (horizontal bar), 5 µm (vertical bar).

## Discussion

Ca²⁺ oscillations are observed in most cell types upon agonist stimulation. Long-term persistent oscillations with regular periodicity were particularly prominent in immune and secretory cells where they regulate secretion, actin dynamics and gene expression (Fewtrell, 1993; Uhlén and Fritz, 2010). Here, by characterizing STIM1 and cortical ER dynamics during receptor-induced Ca²⁺ oscillations in the tumor mast cell model RBL-2H3, we show that STIM1 and ER-PM contact sites exhibit oscillatory dynamics. Importantly, ER-PM contact site oscillations can occur independently of cytoplasmic or ER Ca²⁺ oscillations. This represents a novel example of cortical oscillations involving the SOCE machinery that function upstream of Ca²⁺ oscillations.

The presence of a cortical oscillator that is not merely an effector of Ca²⁺ oscillations has important implications for the mechanisms driving calcium oscillation instability. While early debates considered whether Ca²⁺ oscillations were driven by periodic production of a second messenger from the plasma membrane, the prevailing view is that they are primarily controlled by oscillatory release from the ER store, presumably through CICR. However, the CICR model predicts out-of-phase oscillations between cytoplasmic and ER calcium. Direct experimental confirmation of this prediction using ER-specific calcium sensors alongside cytoplasmic calcium imaging has only been observed in electrically excitable systems, such as neurons and muscle cells (Gu et al., 2024), or in non-excitable cells but under nonphysiological conditions, such as histamine-triggered calcium oscillations in HeLa cells in the absence of extracellular Ca²⁺ (Suzuki et al., 2014).

In RBL cells, Ca²⁺ release from the ER store can sustain a few cycles of cytoplasmic Ca²⁺ pulses under both normal and EGTA-treated conditions, during which ER Ca²⁺ levels exhibit cyclic depletion out of phase with cytoplasmic Ca²⁺ increases. Thus, this mode of oscillatory release-coupled cytoplasmic Ca²⁺ oscillation may correspond to the initial phase of receptor-activated Ca²⁺oscillations. However, in the sustained phase of receptor-activated Ca²⁺ oscillations, ER Ca²⁺ oscillations are absent, suggesting that this phase is distinct from the initial phase of Ca²⁺ spikes. To the best of our knowledge, oscillatory ER Ca²⁺ depletion has not been observed during the sustained phase of receptor-activated Ca²⁺ oscillations in any systems. In exocrine cells, size of agonist-releasable was found to be unchanged when a fresh dose of agonist was added during the interspike period of Ca²⁺ oscillations, indirectly showing that the ER store was full during this period (Shuttleworth and Thompson, 1996). In plant pollen tubes, coupled ER and cytoplasmic Ca²⁺ oscillations have been reported (Resentini et al., 2021), but fluctuations in ER-GCaMP6-210 fluorescence were not attributed exclusively to ER Ca²⁺, as continuous ER membrane movement contributed to the signal—similar to what we observe in mast cells. If ER Ca²⁺ stores remain minimally depleted, ER Ca²⁺ recovery is unlikely to be the rate-limiting step determining oscillation frequency in the persistent phase.

Notably, even in the EGTA phase, ER Ca²⁺ levels were replenished by the time cytoplasmic Ca²⁺ ended,, which was well before the beginning of the next cytoplasmic Ca²⁺ spike, indicating that CICR may not be the primary driver of oscillatory release coupled Ca²⁺ oscillation. This complementary pattern of ER Ca²⁺ depletion and cytoplasmic Ca²⁺ spike was also observed in cardiac myocytes (Brochet et al., 2005).

How these cortical ER and ER-PM contact site oscillations arise is not addressed in the current study. Compared to the mechanisms governing cortical ER recruitment to the plasma membrane, their lifetime and turnover received much less attention. For technical simplicity, most previous studies have employed thapsigargin to deplete ER store which show stable, large-scale redistribution of STIM1 and Orai1/CRACM1 to contact sites (Calloway et al., 2009; Gulyas et al., 2022; Park et al., 2009; Várnai et al., 2007). In one study where physiological stimuli was used, STIM1 oligomerization was induced quickly by receptor activation in RBL cells (Liou et al., 2007), but only the first few minutes after antigen stimulation was investigated, and the dynamics of membrane contact sites during receptor-activated Ca²⁺ oscillations remain largely unexplored. Whether findings from thapsigargin treatment apply to physiological conditions, particularly during the persistent phase of Ca²⁺ oscillations beyond the initial few spikes, remains to be characterized.

Various optogenetic tools have been developed to artificially induce calcium responses by clustering STIM1 cytoplasmic domains (Duan et al., 2017; Kyung et al., 2015; Ma et al., 2020; Pham et al., 2011). However, these approaches have only been successful in modulating calcium dynamics on a slow timescale, with the rise phase alone taking 10 seconds to several minutes and a single calcium pulse lasting several minutes or longer. This is significantly slower than receptor-stimulated Ca²⁺ oscillations in mast cells, which exhibit rise times of 1–3 seconds, total durations of approximately 7 seconds, and oscillation periods of 10–30 seconds. Ca²⁺ oscillations trigged by various agonists in other cell types share similar oscillation periods (Berridge et al., 1988). Therefore, these artificial STIM1 translocation methods fail to replicate the circuit dynamics responsible for the rapid kinetics that drive physiological calcium oscillations.

In separate work, we show that cortical ER oscillations are coupled with phosphoinositide oscillations, particularly PI(4)P and PI(4,5)P₂ (Xiong et al., 2022). Notably, ER-PM contact sites appear to play dual roles in phosphoinositide regulation. On one hand, they act as negative regulators of PI(4)P through transporting PI(4)P to ER, where Sac1, a lipid phosphatase, is localized (Chung et al., 2015; Filseck et al., 2015; Stefan et al., 2011). At ER-PM contact sites, Sac1 degrades PI(4)P, and massive formation of membrane contacts leads to a reduction in PI(4)P levels (Dickson et al., 2016). In yeast, the loss of ER-PM tether proteins results in a substantial accumulation of PI(4)P on the plasma membrane (Quon et al., 2018; Stefan et al., 2011). On the other hand, ER-PM contact sites are essential for the synthesis of PI(4,5)P₂, which replenishes plasma membrane PIP₂ following receptor stimulation (Chang et al., 2013; Chen et al., 2017). The paradoxical role of membrane contact sites in phosphoinositide regulation, along with their mutual feedback mechanisms, requires further investigation. Understanding these dynamics may provide mechanistic insight into how membrane contact site dynamics fine tune dynamical characteristics of Ca²⁺ oscillations. Additionally, it may explain why oscillatory processes were selected as a regulatory strategy to balance competing needs.

## Materials and methods

### Plasmids and reagents

The following reagents were purchased from commercial sources: mouse monoclonal anti-dinitrophenol (DNP) IgE and thapsigargin were bought from Sigma-Aldrich (USA). DNP conjugated bovine serum albumin (DNP-BSA) was from Invitrogen (USA). Constructs of the following plasmids were kind gifts: STIM1-RFP from David Allan Holowka (Cornell University); mEGFP-ER-5a from Michael Davidson (addgene #56455); mCherry-CRY2-5-ptase_INPP5E_, from Pietro De Camilli (Yale University); GFP-Sac1 from Peter Mayinger (Oregon Health and Science University). GCaMP3 (addgene #22692), R-GECO1 (addgene #32444), mCherry-Sec61b (addgene #90994), GFP-MAPPER (addgene #117721), G-CEPIA1er (addgene #58215), CIBN (deltaNLS)-pmGFP (shown as CIBN-GFP-CAAX in text)(addgene #26867), Lyn11-CIBN-GFP (addgene #79572), iRFP (addgene #31857), pmiRFP670-N1 (#79987) from Addgene.

For generating STIM1-Cry2-miRFP670, STIM1 was moved into iRFP using Xho1 and EcoR1 to obtain STIM1-iRFP. Cry2 was obtained by PCR from mCherry-Cry2-ptase_INPP5E_. The PCR product was then inserted in between STIM1 and piRFP using Sal1 and Sac2. pmiRFP670-N1 was digested with Sac2 and Not1 and used to replace iRFP to obtain STIM1-Cry2-miRFP670. For generating miRFP670-Cry2-Sac_TM_, C-termini fragment of hSac1 was first obtained from GFP-Sac1. A separate PCR reaction was performed on vector backbone piRFP-Cry2. PCR product for the C-termini fragment of hSac1 was then ligated into the PCR product of piRFP-Cry2 to obtain iRFP-Cry2-Sac1 Cterminus. Then, hSac1L TM (transmembrane domain) was obtained from iRFP-Cry2-Sac1 Cterminus. A separate PCR reaction was performed on vector backbone iRFP-Cry2-Sac1 C-terminus by overlap PCR. PCR product for hSac1L TM was then ligated into the PCR product of miRFP670-Cry2 vector. Next, miRFP670-Cry2-Sac_TM_ (miRFP670-Cry2-Sac1 LLSTM) was subcloned from miRFP670-Cry2-Sac1 TM by adding a linker between Cry2 and Sac1_TM_ via overlap PCR.

### Cell culture and transfection

RBL-2H3 cells (ATCC, CRL-2256) were maintained in monolayer culture in MEM (Invitrogen) supplemented with 20% fetal bovine serum (Sigma-Aldrich), and harvested with TrypLE Express (Invitrogen). For transient transfections, 2×10^6^ cells were collected and resuspend in 10μl R buffer (Invitrogen), 1μg DNA of each plasmid construct were added and electroporated using the Neon transfection system (Invitrogen) following the manufacturer’s instructions (1200 volts, 20-ms pulse width for 2 pulses). After transfection, cells were spitted into two 35mm glass bottom culture dishes (MatTek) (theoretically 1×10^6^ cells in each dish) and sensitized overnight with 0.4μg/mL anti-DNP IgE. Stable cell line of GCaMP3 was generated by transfection of RBL-2H3 cells for 24 h, then selected in MEM with 0.5 mg/ml G418 (Invitrogen) and sorted for fluorescence. A low expression GCaMP3 stable cell line was used in Fig.2, Fig. 2-Supplemental figure 1 and Fig. 3-Supplemental figure 1.

Four protocols were used for stimulation. In the first protocol (Protocol A, Protocol A-C was according to definition here (Smith et al., 1996)), cells were washed three times with Tyrodes solution (20mM HEPES-NaOH pH 7.4, 135mM NaCl, 5.0mM KCl, 1.8mM CaCl_2_, 1.0mM MgCl_2_ and 5.6mM glucose) before imaging. For antigen stimulation, DNP-BSA was diluted to a final concentration of 80ng/mL in the appropriate buffer and added to anti-DNP IgE sensitized cells. The time antigen is added is stated in the specific figure legends. In the second protocol (Protocol B for Fig. 5), cells were washed with calcium-free buffer (20mM HEPES-NaOH pH 7.4, 135mM NaCl, 5.0mM KCl, 1.0mM MgCl_2_, 5.6mM glucose and 0.5 mM EGTA), and stimulated with antigen in calcium-free buffer. In the third protocol (Protocol C for Fig. 3-4, Fig. 3-Supplemental figure 1, Fig. 4-Supplemental figure 1-2), cells were washed with calcium-free buffer, stimulated with antigen in calcium-free buffer and exchange to calcium-containing buffer. In the fourth protocol, cells were stimulated with 2*μ*M or 10*μ*M thapsigargin.

### Perfusion experiments

In some experiment (Fig. 3-Supplemental figure 1), perfusion system was used. RBL-2H3 cells with GCaMP3 stable expression was cultured on No.1.5, 20mm diameter, round glass coverslips (Marienfeld-Superior) overnight. The coverslips are then transferred to a customized perfusion chamber to form the bottom of the chamber (Chamlide, Live Cell Instrument). A multi-valve perfusion control system (MPS-8, Live Cell Instrument) was used to switch rapidly between prewarmed, calcium-containing Tyrodes buffer or calcium-free buffer.

### Imaging

TIRFM was performed using a Nikon Ti inverted microscope with 3 laser lines (491nm, 561nm, 642nm) at 37°C. The microscope was equipped with an iLAS2 motorized TIRF illuminator (Roper Scientific). TIRFM images were acquired through Nikon objective (Apo TIRF 60X, N.A. 1.49 oil), a quad-bandpass filter (Di01-R405/488/561/635, Semrock) and a Prime 95B CMOS camera (Photometrics). The microscope, camera and illuminator were controlled by Metamorph 7.8 software (Universal Imaging). Confocal images (Fig. 1-Supplemental figure 1) were acquired on a PerkinElmer UltraView Vox Spinning Disk confocal microscope, equipped with a Hamamatsu Electron Multiplying Charge-Coupled Devices camera C9100-50, a 100x Oil objective (Olympus UPlan SApO N.A. 1.4), and controlled by Volocity software. Cells were incubated in a stage-top heated incubator (Live Cell Instrument) set at 37°C and were typically imaged at 2sec per frame for live imaging movies.

### Optogenetics experiment

For optogenetics experiments, we followed our previously published methods (Xiong et al., 2016). Membrane-anchored probe CIBN-GFP-CAAX or Lyn11-CIBN-GFP was used to recruit its cytosolic binding partner Cry2 fused with target proteins upon 491nm light activation. For all the constructs, STIM1-Cry2-miRFP670 and miRFP670-Cry2-Sac1_TM_, 0.1mW of 491nm laser power was used for maximum recruitment. A single or a train of pulses with intervals varying between 20sec to 1min was used to achieve instant or sustained Cry2 recruitment, respectively. We noticed that the calcium sensor GECO1 undergoes reversible photoactivation when stimulated by 491nm laser at powers higher than 0.2mW, therefore we restricted 491nm laser power to 0.1mW for optogenetics to prevent photoactivation of GECO1. For optogenetics on calcium oscillations, we optimized the experimental conditions in order to obtain uniform and robust calcium oscillations. Only RBL-2H3 cells with passage number less than eight were used. We observed that transient transfection of three plasmids together [CIBN-GFP-CAAX (1 µg), GECO1 (1 µg) and STIM1-Cry2-miRFP670 (3 µg)], [Lyn11-CIBN-GFP (1µg), GECO1 (1µg) and STIM1-Cry2-miRFP670 (3 µg)] or [CIBN-GFP-CAAX (1 µg), GECO1 (1 µg) and miRFP670-Cry2-TM_Sac1_ (3 µg)] induce more cell death compared to transfection of two plasmids. To obtain optimal density and higher cell number for imaging, 2 to 3×10^6^ cells were seeded in one 35mm glass bottom culture dishes (MatTek) after transient transfection (theoretically 2-3 higher density than single or double transfection).

### Imaging analysis and statistics

Post-acquisition imaging analysis was performed using Fiji and Matlab (Mathworks). A “reslice” tool and “median” projection filter in Fiji were used to generate kymographs. A “montage” tool was used to generate montages of selected region of interest (ROIs). Pseudo color images in Fig. 1b was generated using “Thermal”, montage in Fig. 1d and kymograph in Fig. 6e was generated using “Gem” lookup table in Fiji. Intensity profiles, wavelet analysis and fast Fourier transform function (FFT) were performed using Matlab. For analyzing the amplitude of fluorescence intensity in Fig. 1f, Fig. 1-Supplemental figure. 3 and Fig. 2g, average intensity of region of interest (30×30 pixels) were quantified by normalization after background subtraction (Ft_cell_-F0_background_)/(F0_cell_-F0_background_). For comparison of the increase in fluorescence intensity in the same cell before and after the light or thapsigargin treatment in Fig. 6c and 7c, intensity is quantified by Ft_cell_-F0_cell_, and paired two-tailed Student *t*-test was used. Peaks before and after optogenetics is compared by Welch’s *t*-test, average data is shown as mean±s.d.

## Acknowledgement

We would like to thank Suet Yin Sarah Fung and Samantha McClellan for calcium imaging, Xiang Le (Jerry) Chua and Su Guo for assistance with molecular biology, Qiang Yuan and Zhifu Wang for assistance with video editing, Bohong Cai for assistance with graphic illustration, XJ Xu for assistance with joy plot. This work was supported by the Singapore Ministry of Education Academic Research Fund Tier 2 (M. Wu, MOE2015-T2-1-122), the Singapore Ministry of Health National Medical Research Council Open Fund Individual Research Grant (M. Wu, NMRC/OFIRG/0038/2017), National Institute of General Medical Sciences of the National Institute of Health Under Award Number R01GM151344 (M. Wu), Natural Science Foundation of China 82301042 (D. Xiong), Sichuan Science and Technology Program Number 2023ZYD0065 and 2023NSFSC0565 (D. Xiong). The content is solely the responsibility of the authors and does not necessary represent the official views of the National Institute of Health.

## Data availability

All data shown and used for quantifications during this study have been provided as source data.

## Materials availability

All materials generated from this study listed in “plasmids and reagents” are available. Requests should be directed to and will be fulfilled by the corresponding authors.

## Code availability

The code used to generate the figures has been previously documented and is openly accessible on GitHub, publicly available at https://github.com/min-wu-lab/ (Tong et al., 2023).

## Author contribution

DX: conceptualization, formal analysis, investigation, writing-original draft, writing-review and editing

CT: formal analysis, investigation

YY: investigation JY: resources

MW: conceptualization, funding acquisition, investigation, supervision, writing-original draft, writing-review and editing

---

**Figure 1-Supplemental Figure 1.**
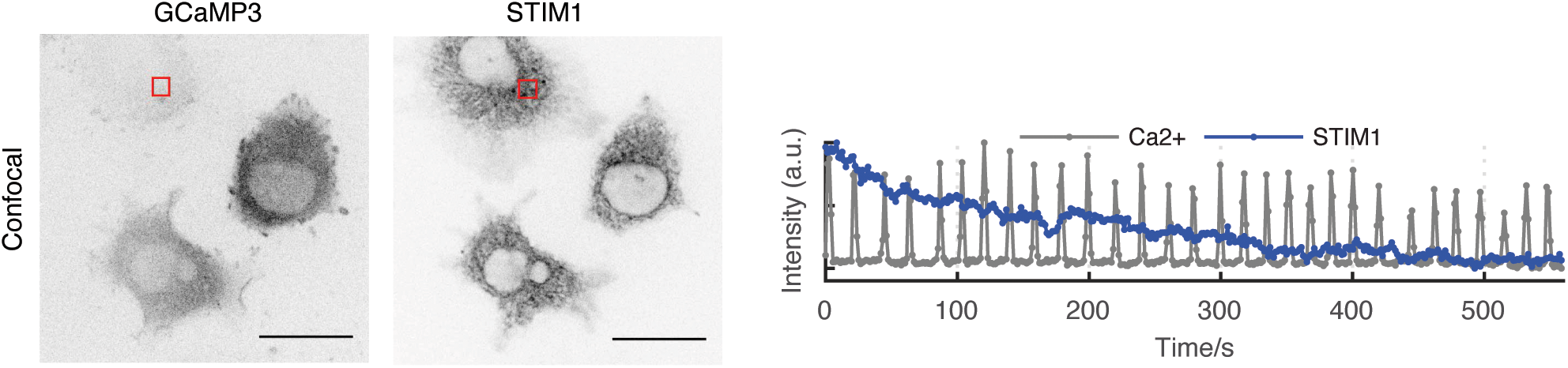
STIM1 oscillations could not be observed by confocal microscopy. Representative single z-plane images and intensity profile of STIM-RFP and GCaMP3 after antigen stimulation, at indicated region of interest on the confocal images (11 cells from 3 experiments). Scale bar = 30 µm.

**Figure 1-Supplemental Figure 2.**
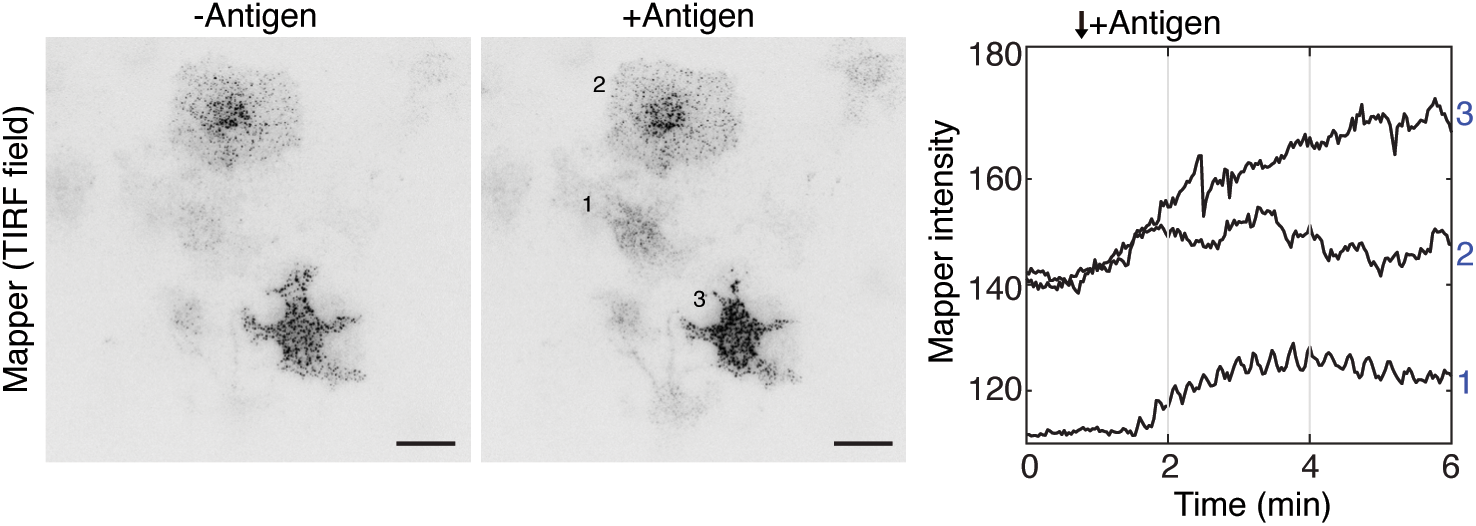
ER-PM contact site oscillations by MAPPER could not be observed in cells with high level of overexpression. TIRF images and intensity profile showing cells expressing different levels of GFP-MAPPER before (-Antigen) and after (+Antigen) DNP-BSA stimulation (10 cells from 3 experiments). Scale bar = 30 µm.

**Figure 1-Supplemental Figure 3.**
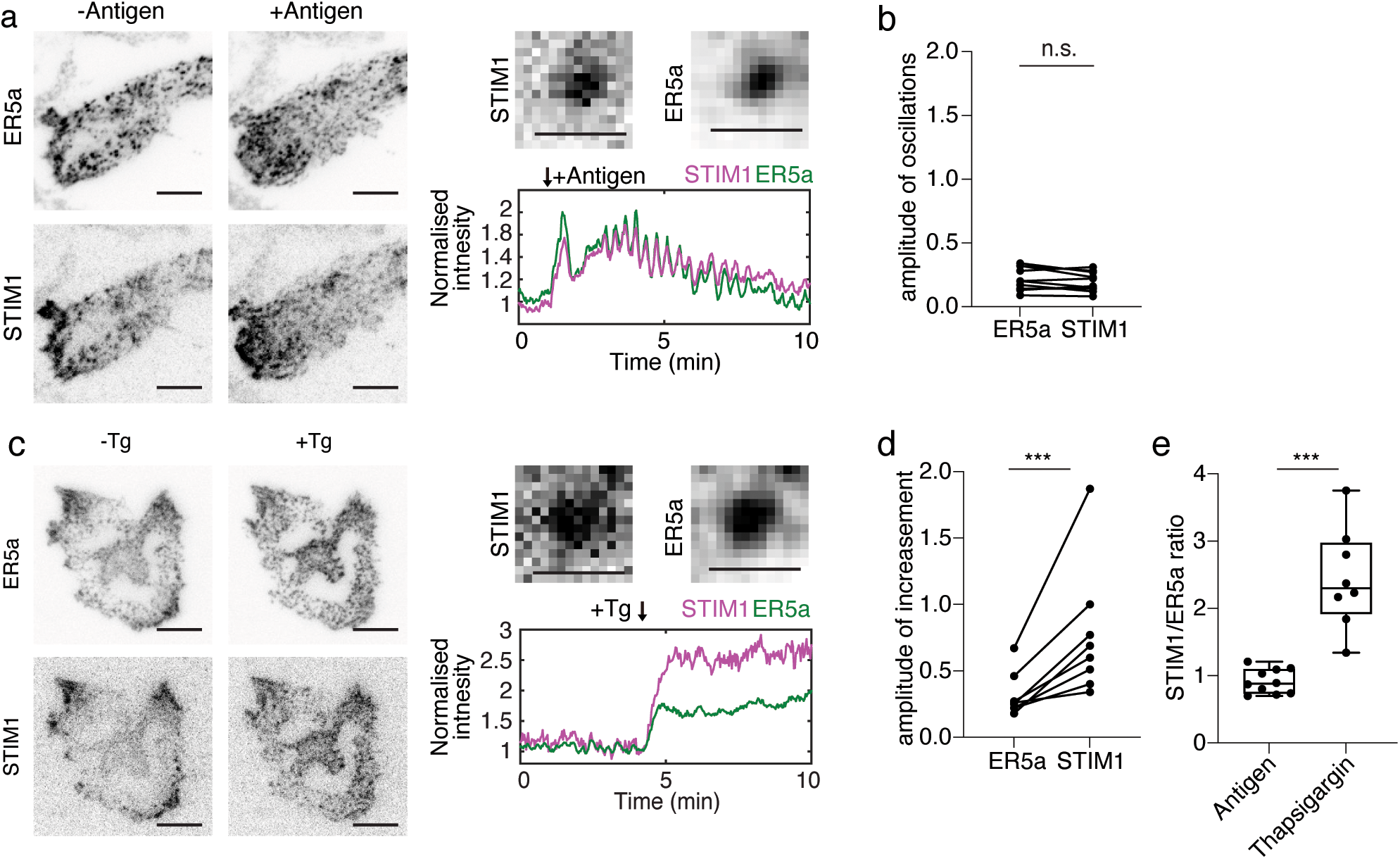
STIM1 and display similar amplitude changes upon antigen stimulation as ER marker ER5a but significantly higher recruitment during Ca²⁺ store depletion. (**a**) Representative TIRF images, zoom-in single punctum images, and normalized intensity profile of cell expressing mEGFP-ER5a and STIM1-RFP upon DNP-BSA antigen stimulation. (**b**) Quantification of the amplitude of oscillation in (a) (n= 10 cells from 3 experiments). (**c**) Representative TIRF images, zoom-in single punctum images, and normalized intensity profile of cell expressing STIM1-RFP and mEGFP-ER-5a upon 2 µM thapsigargin (n=8 cells from 4 experiments). (**d**) Quantification of the amplitude of increase of fluorescence in (c) (n= 8 cells from 4 experiments). (**e**) Ratio of STIM1/ER5a signal upon DNP-BSA antigen stimulation and thapsigargin (8 cells for each condition). (b) and (d) were quantified by paired t-test, (e) was quantified by Welch’s t-test. ***, p<0.001; n.s., not significant. For TIRF images, scale bar= 10 µm; for single punctum images, scale bar=1 µm.

**Figure 2-Supplemental Figure 1.**
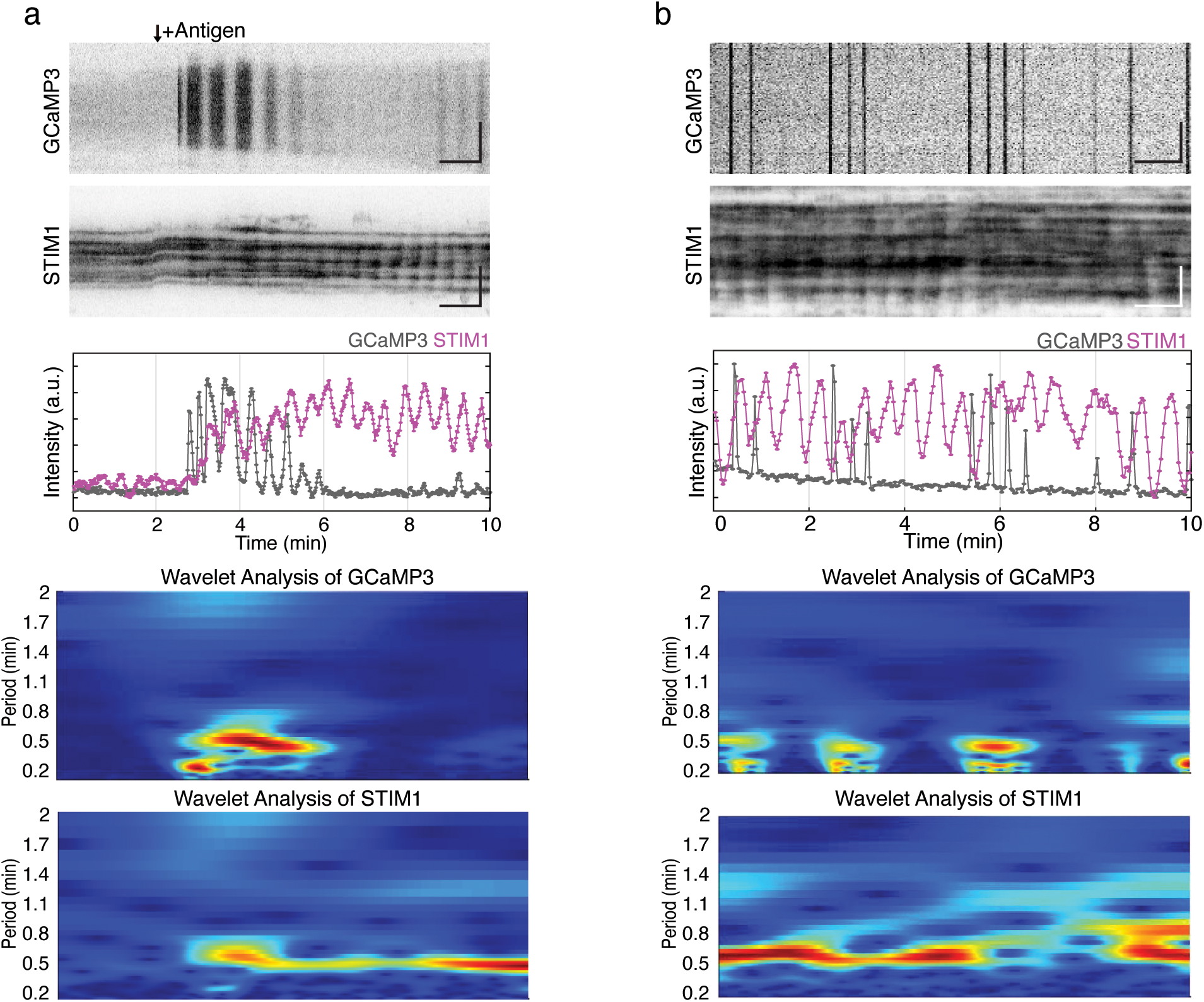
Interconversion between Ca^2+^ oscillation-coupled and uncoupled STIM1 oscillations. (a-b) Kymographs, intensity profiles and wavelet analysis of a cell expressing GCaMP3 and STIM1-RFP showing conversion of Ca^2+^ oscillation-coupled and uncoupled STIM1 oscillation in the initial phase of antigen stimulation (a), or after antigen stimulation during intermittent Ca^2+^ oscillations (b) (n=19 cells from 4 experiments). Scale bar=1 min (horizontal bar), 5 µm (vertical bar).

**Figure 3-Supplemental Figure 1.**
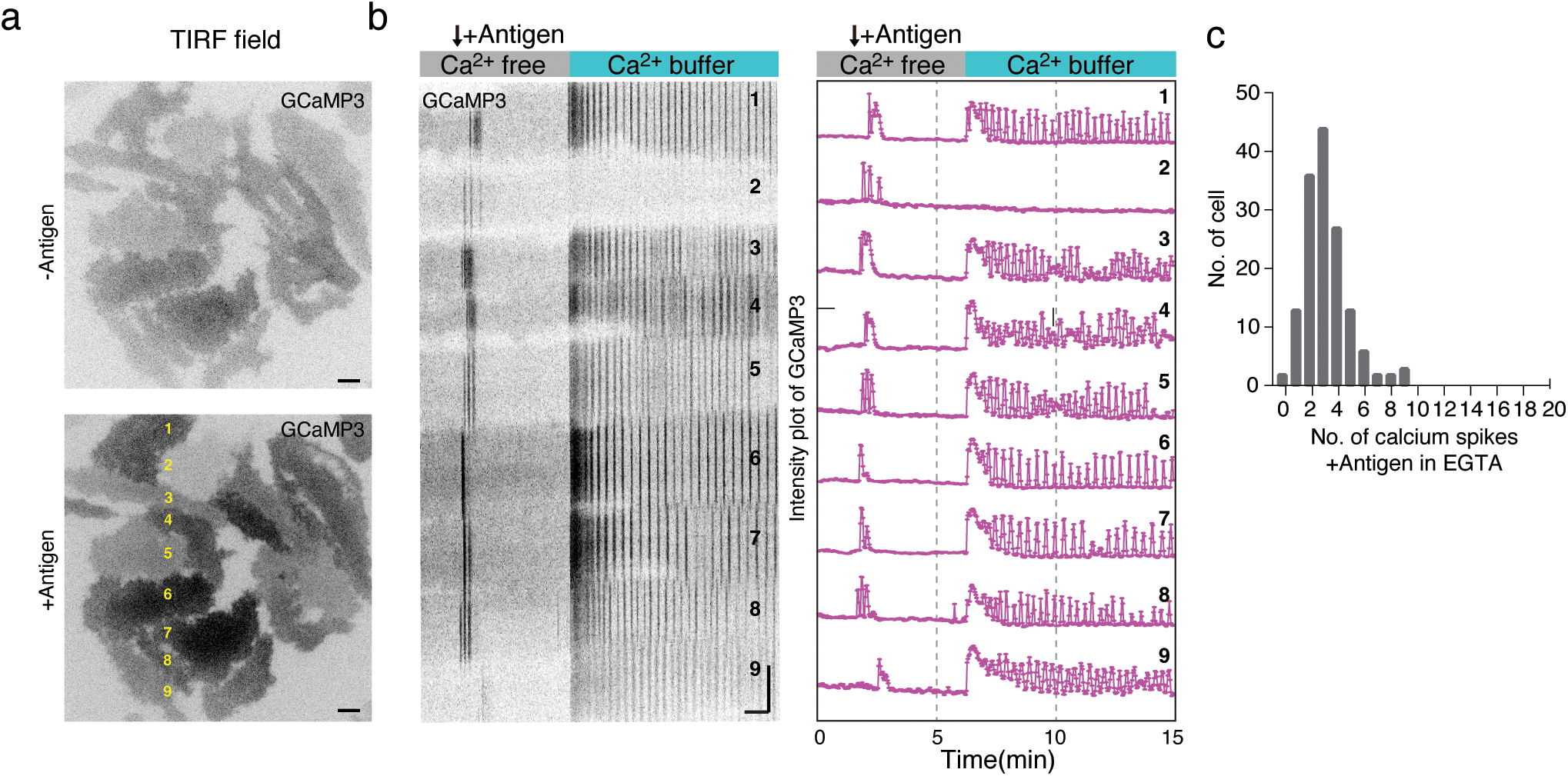
Ca^2+^ oscillations in the absence of extracellular Ca^2+^ oscillation are transient. (a) TIRF images of RBL-2H3 cells stably expressing GCaMP3 stimulated by antigen DNP-BSA in Ca^2+^-free buffer with 0.5mM EGTA first, then exchanged to Ca^2+^-containing buffer (1.8mM Ca^2+^). (b) Kymographs and time-series profiles of the cells marked in yellow in (a). (c) Histogram showing the number of Ca^2+^ spikes upon antigen stimulation in the absence of extracellular Ca^2+^ (n=148 cells from 6 experiments). For TIRF images, scale bar=10µm; for kymographs, scale bar=1min (horizontal bar), 10µm (vertical bar).

**Figure 4-Supplemental Figure 1.**
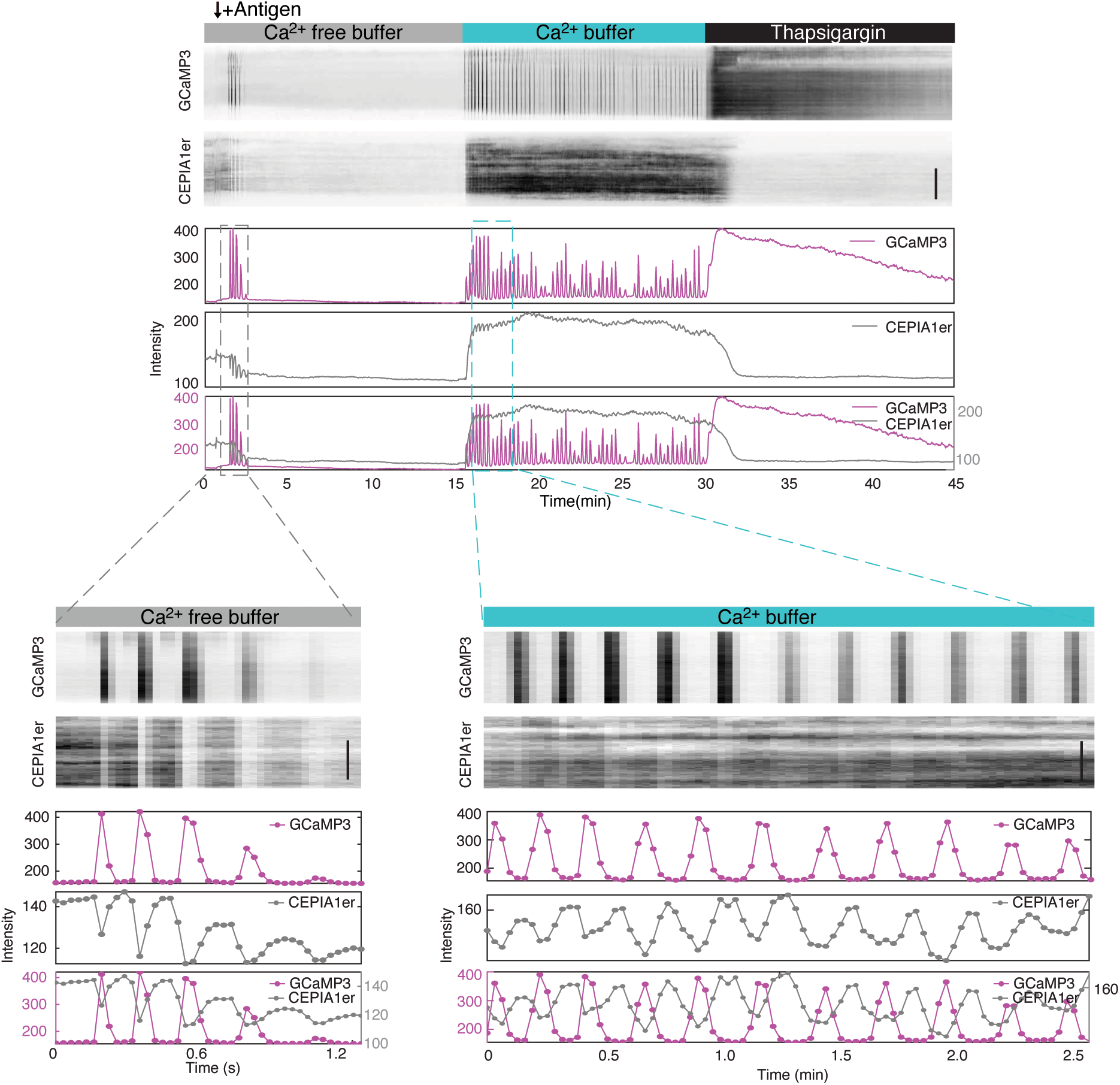
CEPIA1er refills by the time Ca^2+^ spikes ended. Kymograph and intensity profiles of GCaMP3 and CEPIA1er, imaged over 45 minutes in Ca^2+^ free buffer and stimulated with DNP-BSA, followed by replacement with Ca^2+^ buffer and final addition of thapsigargin. Scale bar =10 µm.

**Figure 4-Supplemental Figure 2.**
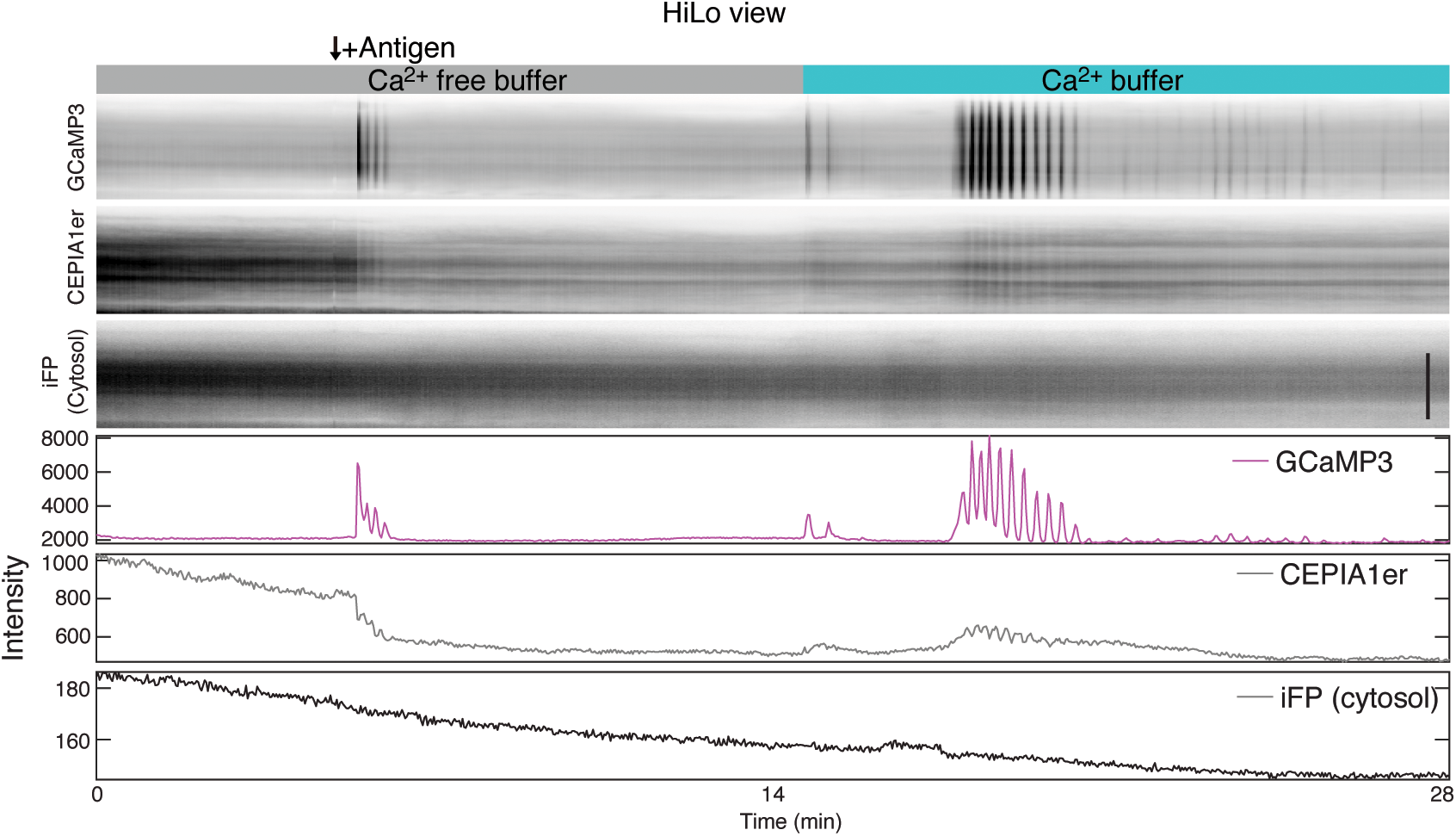
Synchronized reduction in CEPIA1er intensity could be observed at non-TIRF condition when ER movement would not cause oscillations. Kymograph and intensity profiles of GCaMP3 and CEPIA1er, imaged by HiLo over 45 minutes in Ca^2+^ free buffer and stimulated with DNP-BSA, followed by replacement with Ca^2+^ buffer and final addition of thapsigargin. Significant photobleaching occurred during non-TIRF imaging condition. Scale bar = 10 µm.

**Figure 6-Supplemental Figure 1.**
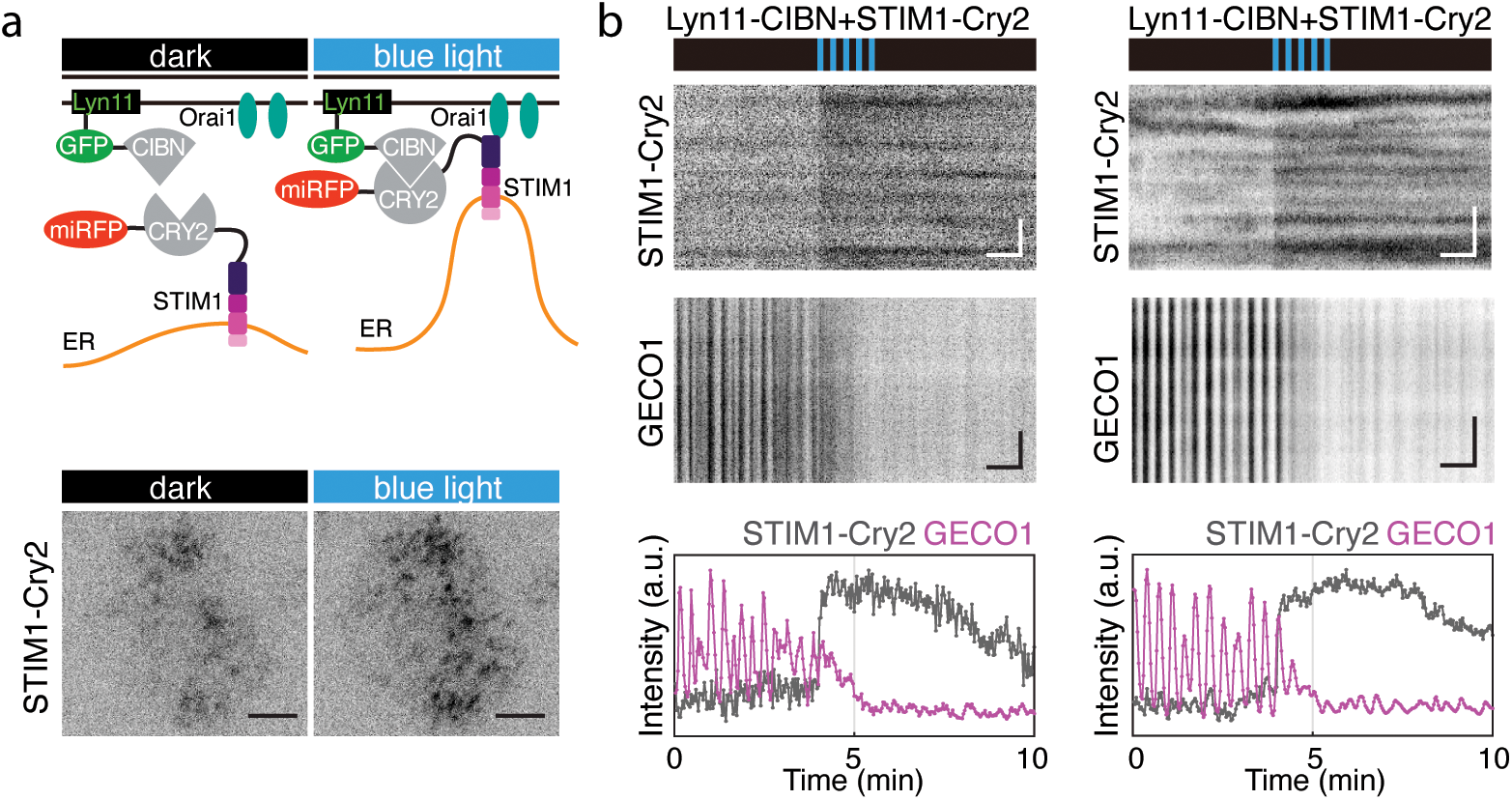
Inhibition of Ca^2+^ oscillations by optogenetically induced translocation of STIM1 does not depend on lipid domain targeting method used. (a) *Top*: Schematic of light-induced plasma membrane translocation of STIM1-Cry2-miRFP670 by Lyn11-CIBN-GFP. *Bottom*: Representative TIRF images of a cell before and after blue light exposure without antigen stimulation (n=20 cells from 4 experiments). (b) Kymographs and intensity profile of STIM1-Cry2-miRFP670 and GECO1 during exposure to blue light pulses. Movies were acquired 10-30min after antigen stimulation. n=15 cells from 4 experiments. For TIRF images, scale bar =10 µm; for kymographs, scale bar =1min (horizontal bar), 5 µm (vertical bar).

**Supplemental video** 1. TIRF movie of cell expressing STIM1-RFP showing cyclic oscillations upon DNP-BSA antigen stimulation near the plasma membrane. Video is acquired at 0.5 sec interval and played at 60x real time. Scale bar=10 µm.

**Supplemental video** 2. TIRF movie of cell expressing GFP-MAPPER showing cyclic oscillations upon DNP-BSA antigen stimulation near plasma membrane. Video is acquired at 2 sec interval and played at 20x real time. Scale bar=10 µm.

**Supplemental video** 3. TIRF movies of cells expressing GFP-MAPPER and mCherry-Sec61β showing coupled oscillations upon DNP-BSA antigen stimulation near plasma membrane. Video is acquired at 2 sec interval and played at 30x real time. Intensity profiles of selected regions of interest are shown accordingly. Scale bar=10 µm.

**Supplemental video** 4. Cyclic STIM1 translocation with or without corresponding calcium signals. Two-color TIRF movie of cell expressing GCaMP3 and STIM1-RFP showing (A) cyclic SITM1 translocation with calcium oscillations, GCaMP3 in grayscale or pseudocolor, and STIM1-RFP in pseudocolor; (B) cyclic STIM1 translocation with irregular calcium spikes; (C) cyclic STIM1 translocation without calcium spikes. Videos are acquired 10-40 min after DNP-BSA antigen stimulation at 2 sec interval and played at 30x real time. Intensity profiles of selected regions of interest are shown accordingly. Scale bar=10 µm.

**Supplemental video** 5. Calcium oscillations are dependent on extracellular calcium. TIRF movie of cell stably expressing GCaMP3 pre-incubated in EGTA containing buffer, stimulated with DNP-BSA antigen stimulation, and change to calcium containing buffer. The video is acquired at 2 sec interval and played at 30x real time. Scale bar=10 µm.

**Supplemental video** 6. TIRF movie of cell expressing STIM1-Cry2-miRFP670 showing the plasma membrane translocation upon 10*μ*M thapsigargin. The movie is acquired without antigen stimulation at 2sec interval and played at 30x real time. Scale bar=10 µm.

**Supplemental video** 7. Two-color TIRF movie of cell expressing CIBN-GFP (not imaged), STIM1-Cry2-miRFP670, and STIM1-RFP showing optogenetic recruitment of STIM1-Cry2-miRFP670 cause coupled STIM1 translocation. Movies are acquired without antigen stimulation at 2sec interval and played at 30x real time. Scale bar=10 µm.

